# Dynamic intestinal stem cell plasticity and lineage remodeling by a nutritional environment relevant to human risk for tumorigenesis

**DOI:** 10.1101/2022.03.17.484800

**Authors:** Jiahn Choi, Xusheng Zhang, Wenge Li, Michele Houston, Karina Peregrina, Robert Dubin, Kenny Ye, Leonard H. Augenlicht

## Abstract

NWD1, a purified rodent diet establishing mouse exposure to key nutrients recapitulating levels that increase risk for intestinal cancer, reproducibly causes *sporadic* intestinal and colonic tumors in the mouse, reflecting human etiology, incidence, frequency and lag with developmental age. Complex NWD1 reprogramming of stem cells and lineages was deconvolved by bulk and scRNAseq, scATACseq, functional genomics and imaging. NWD1 extensively, rapidly, and reversibly reprogrammed Lgr5^hi^ stem cells, epigenetically down-regulating *Ppargc1a* expression, altering mitochondrial structure and function. This suppressed Lgr5^hi^ stem cell functions and developmental maturation of Lgr5^hi^ cell progeny as cells progressed through progenitor cell compartments, recapitulated by *Ppargc1a* genetic inactivation in Lgr5^hi^ cells *in vivo*. Mobilized *Bmi1+, Ascl2^hi^* cells adapted lineages to the nutritional environment and elevated antigen processing and presentation pathways, especially in mature enterocytes, causing chronic, pro-tumorigenic low-level inflammation. There were multiple parallels between NWD1 remodeling of stem cells and lineages with pathogenic mechanisms in human inflammatory bowel disease, also pro-tumorigenic. Moreover, the shift to alternate stem cells reflects that the balance between Lg5 positive and negative stem cells in supporting human colon tumors is determined by environmental influences. Stem cell and lineage plasticity in response to nutrients supports historic concepts of homeostasis as a continual adaptation to environment, with the human mucosa likely in constant flux in response to changing nutrient exposures. Thus, although oncogenic mutations provide a competitive advantage to intestinal epithelial cells in clonal expansion, the competition is on a playing field dynamically sculpted by the nutritional environment, influencing which cells dominate in mucosal maintenance and tumorigenesis.

## Introduction

Plasticity of intestinal cell function underlies seemingly rigid mucosal structural and functional organization. For example, following damage or targeted ablation of Lgr5^hi^ intestinal stem cells (ISCs), multiple cell types can dedifferentiate to function as stem-like cells, interpreted as a damage response to restore homeostasis (1, 2). However, original concepts of homeostasis, by Claude Bernard in the 1860’s as the “milieu interieur”, and Walter Thompson in the 1940’s in coining the term, defined homeostasis as dynamic continual tissue adaptation to its environment, not a single target state returned to following disruption (3). Indeed, we reported that plasticity of intestinal cells to function as stem cells is determined by dietary patterns (4–6) and a recent report documents rapid mucosal adaptation to a high fat chow diet by upregulating fat metabolism pathways (7). Thus, response to nutrients is complex, involving not only adaptive lineage reprogramming, but a major impact on which cells function as stem cells.

The dietary induced stem cell plasticity was in response to NWD1, a purified diet that uniquely causes sporadic intestinal and colon tumors, defined broadly as tumors without clear genetic inheritance and without exposure to known carcinogens. These comprise the majority of human intestinal tumors and are strongly linked to human dietary exposures (4,6,8,9). Moreover, in this model, stem cell reprogramming by NWD1 is a function of nutrient interactions (6), of major importance since scarcity of dietary studies compared to studies of individual nutrients has been emphasized, and it is diet overall that influences cell physiology, disease risk, and progression (10–12). This is not surprising since a vast literature establishes that even strongly penetrant onco- and tumor suppressor genes are always context dependent, with very different effects in different tissues.

There are many strategies for modeling the complex and variable human diet. In the NWD1 model, the principle of nutrient density to adjusts multiple key nutrients to recapitulate exposure levels of each linked to elevated risk for colon cancer in broad segments of high-risk western populations (13, 14). Fat is at 25%, reflecting levels characterizing human high risk, rather than 60% that causes metabolic alterations differing from those caused by level better reflecting human exposure (15, 16). NWD1 also is lower in calcium, vitamin D, methyl donors and fiber, key risk factors for human colon cancer (17). This combination of nutrient risk factors accelerates and amplifies tumor phenotype in multiple genetically initiated models (18–22), and is unique in reproducibly causing *sporadic* tumors of the small and large intestine of wild-type mice, reflecting the etiology, incidence, frequency and lag with developmental age of human colon cancer, the majority of the disease (14,22–24) and that linked to dietary exposures (25). NWD1 fed mice are not obese, but exhibit reduced energy expenditure without altered food consumption or physical activity, causing glucose intolerance, and increased pro-inflammatory cytokines (26). NWD1 elevates Wnt signaling throughout the small and large intestine (27–29), and expands the proliferative compartment (30, 31), as in humans at elevated tumor risk (32–35). It is striking that in NWD1 fed mice, Lgr5^hi^ ISCs are repressed in supporting mucosal maintenance, show reduced mutation accumulation, and reduced efficiency in producing tumors on introduction of an initiating *Apc* mutation, with these functions increasing for Bmi1+ cells (4,6,8,9). These data anticipate the recent report that both Lg5 positive and negative stem cells support human colon tumor growth, the balance determined by environmental factors (36), emphasizing significance of NWD1 shifting which cells function as stem cells in the mucosa.

Here single cell RNAseq, single cell ATACseq, functional genomics and imaging, with rigorous and novel bioinformatic and statistical approaches, de-convolve complex cellular and molecular remodeling of the mucosa increasing risk for the sporadic tumors caused by NWD1. This establishes major new insight into dietary induced risk for sporadic tumors: epigenetic remodeling of chromatin structure of key genes, including a fundamental role of *Ppargc1a* down regulation in altering mitochondrial structure and function specifically in Lgr5^hi^ ISCs; rapidity and reversibility of reprogramming by dietary shift; identity of the cell population mobilized by diet to maintain the mucosa; and elevation of pathways of antigen processing and presentation linked to pro-tumorigenic inflammation. Remarkably, multiple cell and molecular alterations in NWD1 fed wild-type mice parallel pathogenic mechanisms in human inflammatory bowel disease (IBD) (37–39). Therefore, while oncogenic mutations provide a stem cell with a competitive advantage for clonal expansion (1), this competition is carried out on a playing field sculpted and continually remodeled by the nutritional environment, with a major effect on outcome.

## Results

### NWD1 Impact on Lgr5^hi^ intestinal stem cells (ISCs)

Bulk RNAseq analyzed Lgr5^hi^ cells from *Lgr5^EGFP-creER^* mice fed NWD1 or AIN76A control diet for 3 or 12 months from weaning (**Fig 1A**: Arms 1, 2), or 3 months NWD1 then switched to AIN76A and sacrificed at 12 months (Arm 3 – crossover). Of approximately 8,000 genes expressed in isolated Lgr5^hi^ cells, 164 and 63 increased or decreased, respectively, by feeding NWD1 for 3 months (change ≥50% with *P≤0.01* (**Fig S1A**). This increased >3-fold at 12 months to 551 and 229, but reverted to the more limited perturbation in mice fed NWD1 for 3 months switched back to control AIN76A until 12 months (**Fig S1A**). At the pathway level (GSEA), NWD1 significantly up- and down-regulated 25 and 30 pathways, respectively, in Lgr5^hi^ cells at 3 months, many alterations persisting to 12 months of feeding, and reversed upon shift back to control AIN76A diet (**Fig S1B**). Thus, NWD1 had a major effect on transcriptional profile of Lgr5^hi^ cells.

**Figure 1:**
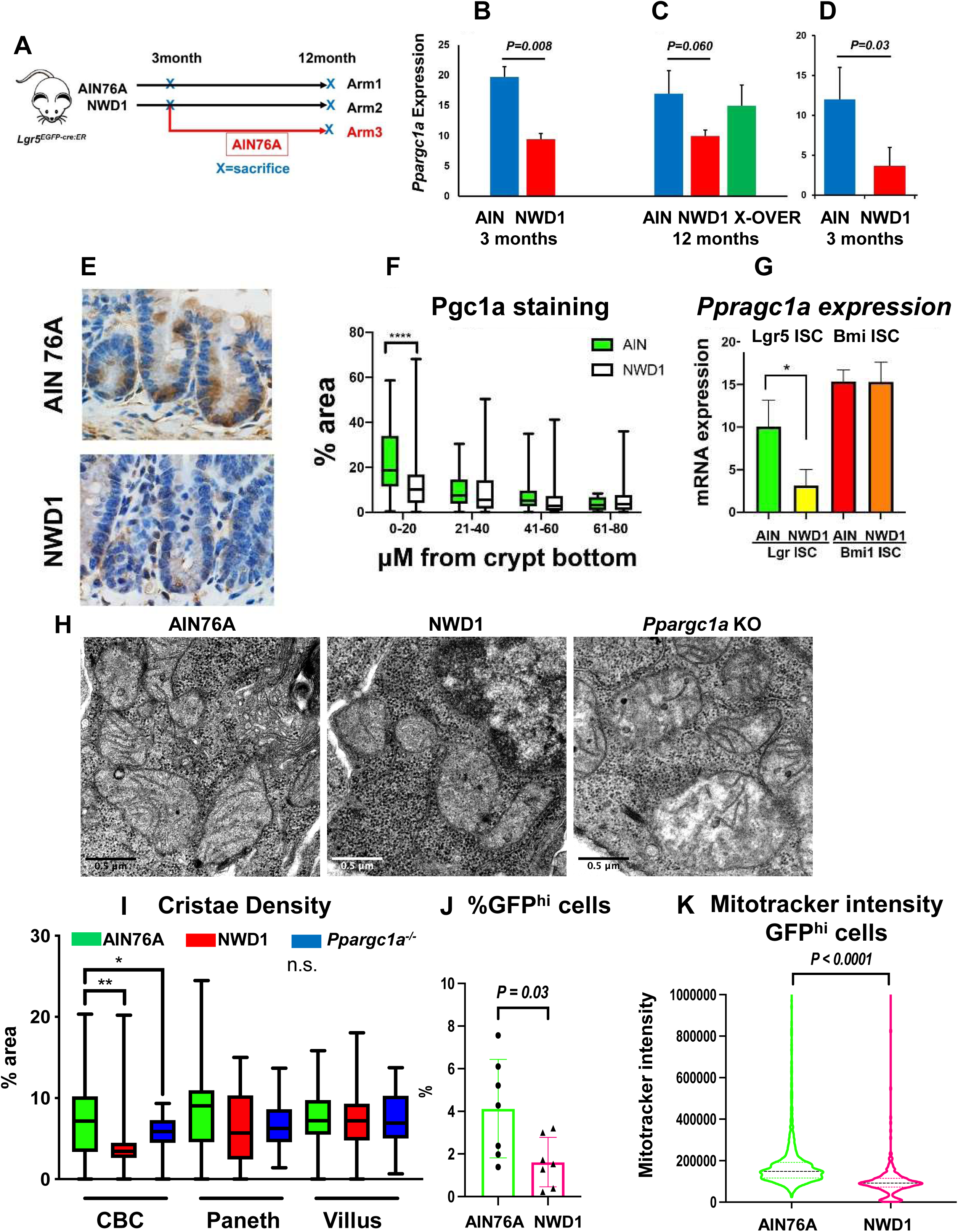
Dietary impact on intestinal stem cells: **A,** *Lgr5^EGFP.cre:ER^* mice fed AIN76A or NWD1 for 3 or 12 months from weaning, or NWD1 for 3 months then switched to AIN76A for 9 months (cross-over); **B - D,** *Ppargc1a* expression by bulk RNAseq of Lgr5^hi^ cells of 2 different mouse cohorts fed either NWD1 or AIN76A for 3 months from weaning (**B, D**), or for NWD1 or AIN76 for 12 months compared to NWD1 for 3 months and then switched to AIN76A for an additional 9 months (**C -** Arm 3, Fig 1A); **E, F** Pgc1a immunohistochemistry and quantitation in crypts of mice fed AIN76A or NWD1 for 3 months; **G**, *Ppargc1a* expression in Lgr5^hi^ and Bmi1+ cells from bulk RNAseq analysis of mice fed different diets for 3 months; **H**, examples of mitochondria in wild-type mice fed AIN76A or NWD1, or mice fed AIN76A and *Ppargc1a* homozygously inactivated in Lgr5 cells (800X); **I**, quantitation of mitochondrial cristae density in CBC, Paneth, and villus cells evaluated at 5000X magnification; **J,** FACs analysis of Lgr5^hi^ cell number; **K**, Mitotracker staining of mice fed AIN76A or NWD1 for 3 months. *(*P<0.05, **0.01, ***0.001*).

The Oxphos and the TCA cycle pathways were down-regulated by NWD1 at 3 months, persisted until 12 months, and reversed by 12 months after switching back to control diet at 3 months (**Fig S1B, S2**), necessary for intestinal *Lgr5^hi^* cells, hematopoietic, and embryonic cells to function as stem cells (40–44). NWD1 down-regulated 58 of 132 genes in the Oxphos pathway at 3 months, confirmed in an independent cohort of mice also fed NWD1 or AIN76A for 3 months (6), with mean Oxphos gene expression decreased by 14% and 11% in the two data sets (**Fig S3A**). Down regulation of most Oxphos genes was modest, but the altered profile was highly significant (*P=10^-26^*, Fisher’s exact test in comparison to the 8,000 member expressed gene set). The down regulated profile and pathway persisted at 12 months (**Fig S1B, S2, S3B**), encompassing multiple genes encoding subunits of the mitochondrial electron transport chain (**Fig S3C).** Greatest down regulation in both cohorts were *Cox6a2* and *Cox7a1* of cytochrome oxidase, the terminal respiratory chain enzyme, and *Sdhb*, and *Sdhd* of complex II, succinic dehydrogenase (**Fig S3A**), down-regulated in CRC and other cancers causing succinate accumulation and tumor development (45, 46). In crossover to control diet (Arm 3), 52 (82%) of down-regulated Oxphos genes increased towards control levels, also highly significant (**Fig S3B;** *P=10^-18^*), reflecting GSEA reversal (**Fig S1B, S2)**.

*Ppargc1a*, encoding Pgc1a, is a master regulator of metabolism and mitochondrial biogenesis (47, 48) and *Ppargc1a* overexpression elevates mitochondrial biogenesis and function of colonic epithelial cells, altering their fate (49). Importantly, feeding NWD1 for 3 months reduced *Ppargc1a* expression in Lgr5^hi^ cells by ≥60% in both mouse cohorts (**Fig 1B,D** *P=0.008 and 0.03*), similarly decreased at 12 months of NWD1 feeding (*P=0.06*), and switchover back to control AIN76A produced levels indistinguishable from mice continuously fed AIN76A from weaning (**Fig 1C**). By immunohistochemistry, the encoded protein, Pgc1a, was highest in crypt cells of control fed mice, but much lower in crypts of NWD1 fed mice, paralleling decreased *Ppargc1a* expression in Lgr5^hi^ stem cells (**Fig 1E**). Quantification across multiple crypts-villi established decreased Pgc1a in crypts of NWD1 fed mice (*P<0.01*), but much lower expression, and no difference with diet, at any position above 20 microns of the crypt base (**Fig 1F**). Further confirmation was decreased *Ppargc1a* expression in Lgr5^hi^ cells, but not in Bmi1+ cells most abundant in cells further up into the villi (**Fig 1G**). Quantitatively, cristae density in crypt base epithelial cells decreased in NWD1 fed mice (*P<01*), with no change in cristae density in Paneth cells at the base, or in cells higher up (**Fig 1H,I**). Finally, GFP^hi^ cells of *Lgr5^EGFP^* mice fed NWD1 decreased by 60% (**Fig 1J**, *P=0.03*) and their retention of mitotracker dye decreased (**Fig 1K**, *P<0.0001*), an index of mitochondrial function. Therefore, NWD1, in lowering *Ppargc1a* expression, its encoded Pgc1a protein, and expression of many genes encoding multiple proteins of the electron transport chain, altered mitochondrial structure and membrane potential in stem cells at the crypt base.

### Mechanistic role of *Ppargc1a* down regulation

*Ppargc1a* was inactivated hetero- or homozygously in Lgr5^hi^ cells of mice fed control diet (*Lgr5^EGFP-cre:ER-^*, *Ppargc1a^loxp/loxp^ ^or^ ^loxp/+^, Rosa26^tom^*mice). 3 days after homozygous inactivation, effects of feeding NWD1 were phenocopied: Lgr5+ cell lineage tracing was significantly suppressed (**Fig 2A,B**, *P=0.02*), and CBC cell cristae density significantly decreased (*P<0.05*) with no alteration in Paneth or in villus cells (**Fig 1H,I**).

**Figure 2:**
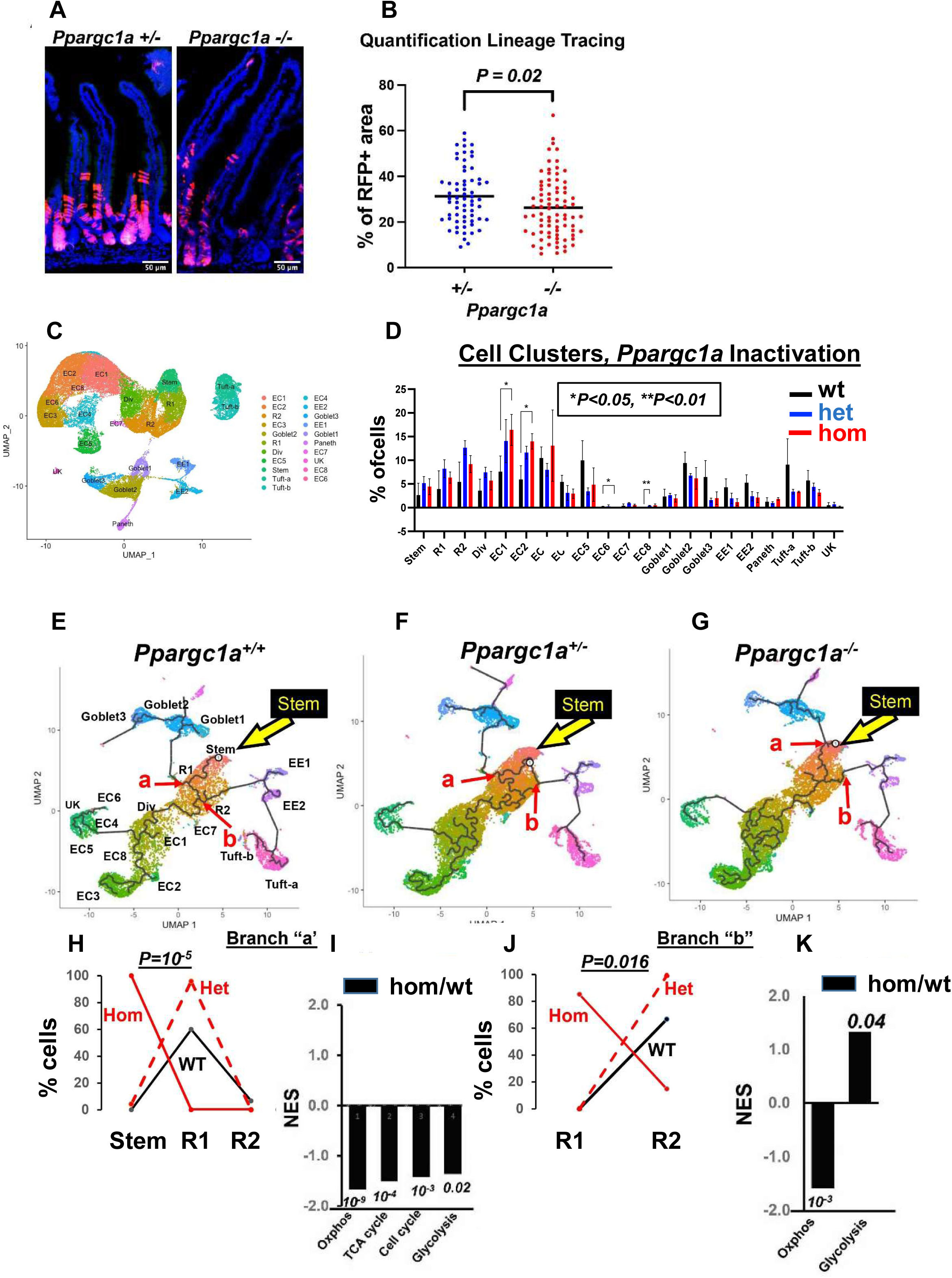
Genetic inactivation of *Ppargc1a* in Lgr5 cells: **A,** suppressed lineage tracing from Lgr5^hi^ cells by homozygous inactivation of *Ppargc1a* in *Lgr5^cre:er-GFP^, Ppargc1a^F/+ or-/-^, Rosa26^tom^* mice fed AIN76A diet, 3 days post a single Tam injection; **B**, quantification of lineage tracing; **C,** cluster map and cell types, N=3 independent mice for each genetic group (9 scRNAseq libraries); **D**, epithelial cell distribution in clusters/lineages in wild-type mice or with heterozygous or homozygous *Ppargc1a* inactivation targeted to Lgr5^hi^ cells; **E-G**, trajectory analysis from scRNAseq of total intestinal epithelial cells from wild-type mice or 3 days after hetero- or homozygous *Ppargc1a* inactivation: yellow arrow, Stem cell cluster; **H, I** cell type distribution at branch point “a” and “b”, respectively (red arrows in E-G) of wild-type, het and hom inactivation of *Ppragc1a* and GSEA of pathways of *Ppargc1a^-/-^* compared to wild-type mice at those branch points (numbers indicate *P values*).

To define how this remodels stem cells and the mucosa, scRNAseq analyzed FACs isolated intestinal epithelial cells (Epcam+, CD45neg) from mice 3 days after *Ppargc1a* inactivation in Lgr5^hi^ cells. 20 clusters were identified and aligned with intestinal cell types and lineages (**Fig 2C** **-** (Methods)). A minor cluster could not be aligned (UK; <1% of cells in any single cell library). Cell proportion in each cluster was similar among genotypes, but with increased early enterocyte populations (EC1 and EC2) by heterozygous inactivation that increased further and was statistically significant with homozygous inactivation (**Fig 2D**). Trajectory analysis placed EC1 proximal to Dividing cells, EC2 emerging on that trajectory, and other EC cells and lineages arising further down-stream (**Fig 2E-G**). This predicted trajectory using Monocle was validated by interrogating markers that identify physical cell position along the crypt-villus axis (**Fig S4A,** (50)). Therefore, decreased EC1 and 2 cell number within 3 days of *Ppargc1a* inactivation in Lgr5^hi^ ISCs was consistent with these cells being the most immediate cells emerging from the progenitor cell compartments.

Repressed mitochondrial function suppresses ability of Lgr5^hi^ cells to function as stem cells (40–44), mitochondrial function determines balance between stem cells retaining self-renewal capacity or differentiating (51), and mitochondrial fusion and hence activation is necessary for intestinal stem cell progeny to differentiate (52). Therefore, impact of *Ppargc1a* inactivation in Lgr5^hi^ cells on cell progression through progenitor cell compartments was determined (**Fig2 E-G**). As expected, progressive maturation of progenitor cells began at the Stem cluster, with maturation of progenitor cells through branch points giving rise to committed lineages. At branch point “a”, developmental trajectory from the Replicating 1 cell compartment (R1) bifurcates to secretory goblet cell lineages or R2 cells (red arrow, **Fig2 E-G**). For both WT and *Ppargc1a* het mice, R1 cells were the major cell type at branch point “a” with few or no Stem cells (**Fig 2H**). Thus, at this early branch point, developmental transition from Stem to R1 had taken place. However, with homozygous *Ppargc1a* inactivation, 100% of cells at “a” were still identified as stem cells, a major, highly significant, repression of re-programming with maturation from Stem to R1 cells (**Fig 2H**; *P=10^-5^*, by a weighted linear regression based on cell number at each cluster for each mouse). The altered cell distribution at branch “a” with homozygous inactivation was characterized by significant suppression of Oxphos, TCA cycle, cell cycle, and glycolysis pathways (**Fig 2I**), reflecting the altered mitochondrial structure and function with *Ppargc1a* homozygous inactivation1 (**Fig 2H,I**).

With further progression along the validated trajectory, branch point “b” was transition from R2 to enteroendocrine and Tuft cells (**Fig 2E-G**). At “b”, most cells in WT or het mice had progressed to the R2 compartment, but with homozygous *Ppargc1a* inactivation, 90% remained in R1 (*P=0.016;* **Fig 2J**), with highly significant repression of the Oxphos pathway persisting (**Fig 2K**). Therefore, homozygous *Ppargc1a* inactivation altered Stem cell maturation to R1 cells, and repressed transition out of R1 with down-regulation of the Oxphos pathway, and a parallel enrichment in the glycolytic pathway. Consistent with retarded cell developmental maturation is the significant increase in EC1 and EC2 early enterocyte populations with homozygous *Ppargc1a* inactivation (**Fig 2D**).

### Rapidity and reversibility of dietary alterations

scRNAseq analyzed intestinal epithelial cells of mice fed AIN76A for 3 months (**Fig 3A**, Arm 1), switched to NWD1 for 4 days (Arm 2), and the 4d NWD1 mice switched back again to AIN76A for 4 days (Arm 3). Stem, replicating, dividing cells and all lineages were identified, with no significant changes in cell distribution for clusters among the 3 arms in these short time periods (**Fig S5A,B**), and cell trajectory confirmed by expression of genes known to be localized to different positions (**Fig S4B**). For the stem cell cluster, 4 days of NWD1 altered 30 pathways with high significance (*P=10^-3^ to 10^-43^*) including repression (negative Z-score) of the TCA cycle and Oxphos (*P=10^-6^ and 10^-43^*, respectively; **Fig 3B,C**). 19 of the 30 pathways reverted to control values in 4d NWD1 mice switched back to AIN76A for 4 days. This included the TCA cycle pathway, down regulated by 4d of NWD1 (*P=10^-6^*), then up-regulated when switched back to AIN76A ((**Fig 3B,C**). Expression of 8 genes encoding TCA cycle enzymes were down-regulated by 4d NWD1 and reversed by 4d switch-back to AIN76A (**Fig 3D**), accounting for the highly significant GSEA for dietary shifts (**Fig 3B.C**). 4d of NWD1 decreased expression of 51 Oxphos pathway genes (**Fig 3E**), consistent with the highly significant negative GSEA (*P =10^-43^*), **Fig 3B,C**), but only 15 reversed on AIN76A switch back (**Fig 3E**), reflecting the absence of reversal by GSEA (**Fig 3C**). Therefore, within 4 days of shift to NWD1, stem cells at the crypt base underwent significant mitochondrial re-programming, in large part, but not entirely, reversed within 4 days of shift back to control diet.

**Figure 3:**
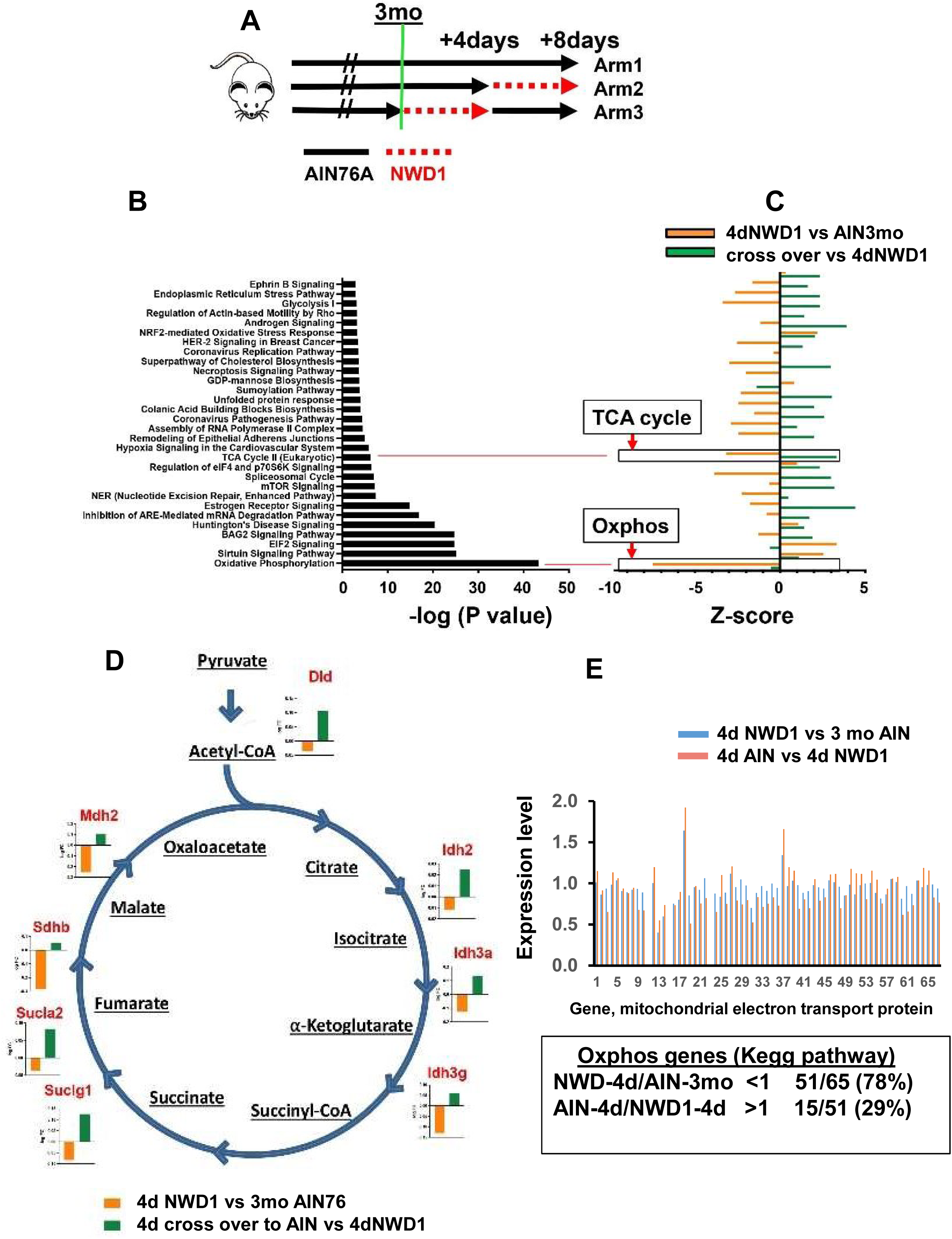
Rapid reprogramming of cells by dietary shift: **A,** mice fed AIN76A for 3 months (Arm1), switched to NWD1 for 4 days (Arm 2), or then switched back to AIN76A for 4 days (Arm 3), total Epcam+, CD45neg epithelial cells FACs isolated and analyzed by scRNAseq; **B,** pathways significantly altered by rapid dietary shifts and their negative log *P* value for significance of pathway change; **C**, magnitude of change of each pathway under the different dietary conditions; **D,** TCA cycle genes repressed by switching mice from AIN76A to NWD1 for 4 days and then elevated when mice switched back to AIN76A control diet for 4 days; **E,** altered expression of each gene in the Oxphos pathway by 4 day shift from AIN76A to NWD1, and response of each to subsequent switch back to AIN76A for 4 days.

### NWD1 mobilization of Bmi1+ cells

NWD1 expands the mouse intestinal proliferative cell compartment (30, 31), recapitulating expansion in humans at elevated CRC risk (32–35), consistent with NWD1 induced lineage tracing from Bmi1+ cells above the crypt base that persists for months as long as mice are fed NWD1 (6, 8). Moreover, in targeting *Apc* inactivation to Bmi1+ cells, tumor development increased when the mice were fed NWD1, paralleling the NWD1 induction of lineage tracing, and hence stem cell-like functions, in these cells (4, 6).

Following injury to Lgr5^hi^ ISCs, stem cell functions can be acquired by dedifferentiation of cells above the crypt base (1, 2). We have emphasized this plasticity also characterizes adaptation of cells to their nutritional environment (4–6). To dissect this in much greater depth, scRNAseq data were generated for Bmi1+ cells and their marked epithelial cell progeny isolated from *Bmi1^cre-ERT2^, Rosa26^tom^* mice fed diets for 3 months from weaning (Tom+, Epcam+, CD45neg cells). The Bmi1 marker was activated at 3 months by a single Tam injection, each mouse continuing on its diet for 3 or 66-70 days before sacrifice to analyze the cells within 3 days of marking, or after 2 months of their accumulation in the mucosa (**Fig 4A; S6A,B**). Stem, replicating, dividing, and all cell lineages were identified with similar cell cluster distributions among cell types regardless of diet (**Fig 4B; Fig S6B,C**). Therefore, despite NWD1 repression of Lgr5^hi^ stem cell functions, the mucosa was maintained. As above, predicted developmental trajectory of Bmi1+ cells was confirmed by expression of markers that identify cells at different positions along the crypt-luminal axis (50) (**Fig S4C**).

**Figure 4:**
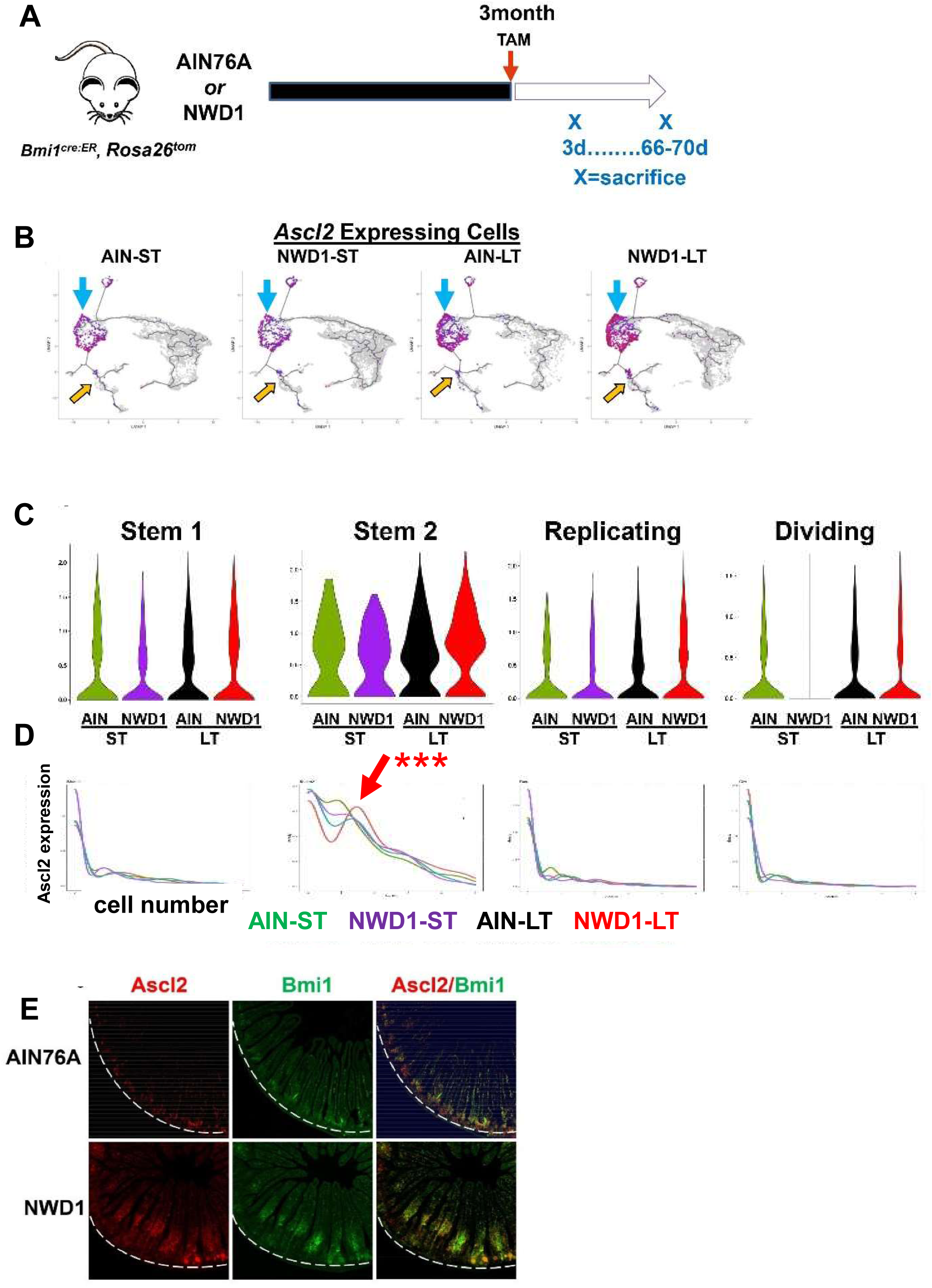
scRNAseq of Bmi1+ intestinal epithelial cells in response to diet: **A,** Rosa26^tom^ marked Epcam+, CD45neg epithelial cells FACs isolated from *Bm1^cre:er^, Rosa26^tom^* mice fed AIN76A or NWD1 for 3 months, then sacrificed at 3 or 66-70 days after TAM injection to activate the Rosa^tom^ marker (shorter, longer term, respectively) and cells analyzed by scRNAseq (N=2 for each group); **B,** cell trajectory analysis as a function of diet and time post marking by Tam injection: Blue arrows, *Ascl2* expressing cells among stem and progenitor cells. or yellow arrows, in goblet and enteroendocrine cells; **C**, **D,** expression of *Ascl2* per cell in Stem1, 2, Replicating and Dividing cells clusters: red arrow in **D** for Stem 2 cells is a population that expressed *Ascl2* at a higher level (***statistical analysis detailed in text); **E,** *Ascl2* and *Bmi1* expression by *in situ* hybridization in mice fed diets for 3 months: white dotted line demarks the crypt base.

Expression of the stem cell transcription factor Ascl2 drives dedifferentiation of Bmi1+ cells to replace damaged Lgr5^hi^ cells (1, 2). Regardless of diet or time post Bmi1+ cell marking, *Ascl2* was expressed in the Bmi1+ Stem1 and 2, Replicating and Dividing cells in trajectory analysis of the data (**Fig 4B**, blue arrows, quantified in **Fig S6D)**. A small number of Tuft and goblet cells also expressed *Ascl2* where goblet and enteroendocrine lineages diverged (**Fig 4B**, orange arrows, and **Fig S6D**).

Among the 4 progenitor cell compartments, mean *Ascl2* expression per cell was highest in Stem2 (**Fig 4C**). *Ascl2* expression level across each individual cell was low per cell for Stem 1, Replicating and Dividing cells, with no significant difference by diet or length of time post Tam cell marking (**Fig 4D**). In contrast, for Stem 2, expression per cell was heterogeneous, with a subset of Stem2 cells expressing elevated *Ascl2* that accumulated longer term (**Red arrow**, **Fig 4D**). This pattern was highly significant by a likelihood ratio test (*P=0.01*, by a negative binomial regression-negative binomial mixed effect model, with regression and fit using R functions glm.nb and glmer.nb, respectively (53), assuming each cell as independent, and number of *Ascl2* reads for each cell as response). Post-hoc analysis by a similar negative binomial regression showed NWD1-LT differed significantly from the other 3 groups in *Ascl2* expression (*P=0.002*). Finally, potential that the difference between NWD1-LT and the other groups was random was tested by a negative binomial mixed effect model with mice per dietary group treated as random effects; the likelihood ratio test was *P = 0.11*, indicating the alternate hypothesis that effects were random is false.

*In situ* hybridization localized *Bmi+*, *Ascl2^hi^*expressing cells confined to the crypt base in AIN76A mice, but in NWD1 mice, elevated *Ascl2* was in Bmi1+ cells above the crypt base (**Fig 4E**), as in mice after acute Lgr5^hi^ cell damage (1, 2), although in response to NWD1, these cells are further above the crypt (Discussion).

### Remodeling of the mucosa by the mobilized Bmi1+ cells

In the validated developmental trajectory of Bmi+ derived cells, there are 2 early trajectory branch points among progenitor cell populations (**Fig 5A**, red arrows 1 and 2). At branch point “1”, Stem1 cells give rise to enteroendocrine, goblet and Paneth cell secretory lineages. Data from libraries from different mice gave identical results, and there were no significant changes in cell type representation by diet at Branch 1 either shorter or longer term after Bmi1+ cell marking (**Fig 5B**). However, at branch point 2 there were highly significant differences: in Bmi1+ cells shorter term after marking (**Fig 5C**, **top**), cells from AIN76A control fed mice were equally divided between R and Div cells, but in NWD1 fed mice, all cells remained in R with none in the Div cell compartment, a highly significant dietary linked difference (*P=0.001*), with replicate identical patterns from independent mice. This became more pronounced in Bmi1+ cells that accumulated longer term (**Fig 5C, bottom**), with nearly all cells at this branch point 2 of AIN76A fed mice Div cells, but in contrast, for NWD1 mice, 100% of the cells still had a transcriptional profile of R cells. Independent mouse replicates were again identical, and the dietary differences highly significant (*P=001*). This novel dietary effect on suppressing developmental progression of cells in their trajectory along the crypt-villus axis was also reported in mice in which genetic manipulation of mitochondrial structure down-regulated stem cell function (52)and in human IBD patients (39). Notably, R cluster cells at branch point 2 longer term after marking express much higher *Ascl2* levels (**Fig 5D**, *P=10^-8^*) similar to the *Bmi1+, Ascl2^hi^* cells that accumulate above the crypt base in NWD1 mice (**Fig 4E**). and the Stem2 subset in NWD1 that express higher *Ascl2* in Bmi1+ cells that accumulate longer term after marking (**Fig 4D**).

**Figure 5:**
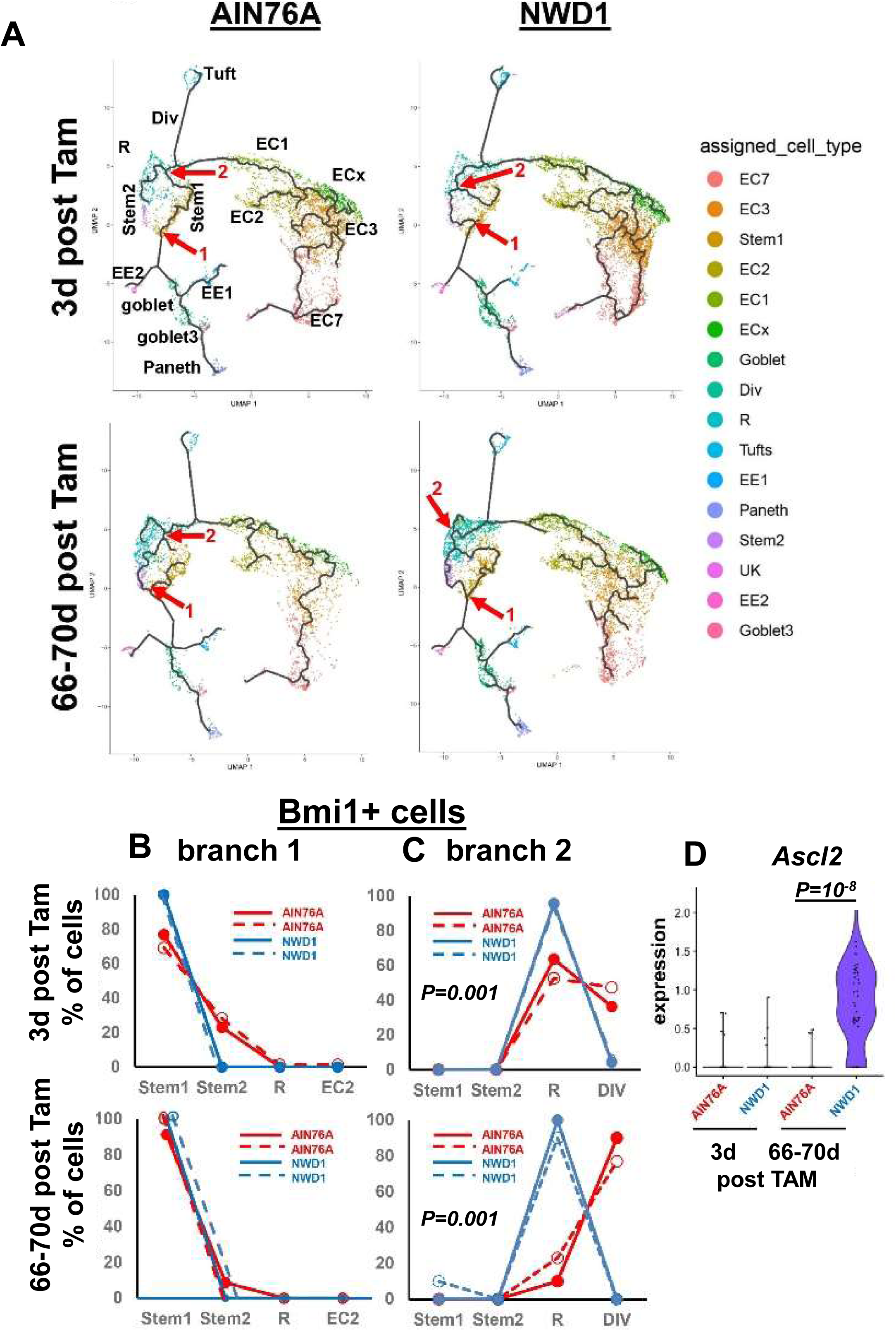
**A**, trajectory analysis of Bmi1+ marked cells at 3d or 66-70 days post Tam activation of the Bmi1 marker, annotated with individual cell types: red arrows/numbers denote branch points analyzed; **B, C** cell type distribution at branch points “1” and “2” (red arrows in A); **D**, *Ascl2* expression per cell at branch point 2.

### NWD1 cell reprogramming is pro-inflammatory and adaptive

Although Bmi1+ marked cells give rise to all lineages at equivalent levels in NWD1 and AIN76A fed mice, NWD1 altered gene expression profiles in all Bmi1+ cell populations both long and short term after marking. This is most pronounced in enterocytes, especially in EC7 enterocytes accumulating longer term (**Fig 6A**). EC7 is predicted to be the most mature enterocyte cell population, validated by markers that identify villus tip cells ((50), **Fig S4C**).

**Figure 6:**
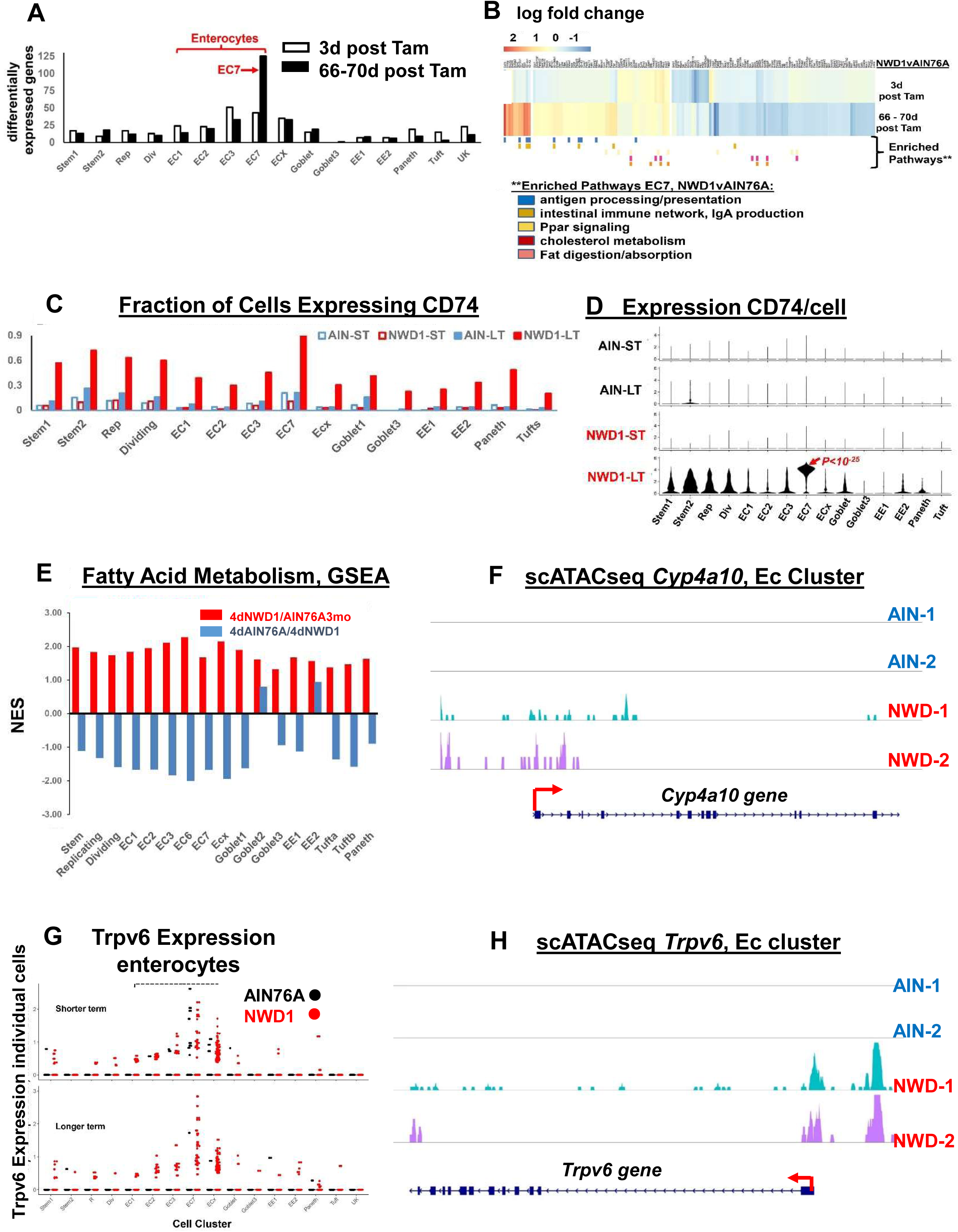
NWD1 reprogramming and adaptation of cells: **A,** number of differentially expressed genes (>50% *and P=0.01*) in Bmi1 cell clusters of Fig S6B; **B,** Heat map of genes differentially expressed by diet in EC7 cells and cell pathways enriched (GSEA) as a function of diet in Bmi1+ EC7 cells longer term post marking of Bmi1+ cells (statistical analysis detailed in text); **C**, fraction of cells in each cluster expressing CD74; **D**, CD74 expression per cell, in each cell cluster as a function of diet and time post marking of Bmi1+ cells; **E,** GSEA analysis for each cell cluster for the fatty acid metabolism pathway in the rapid dietary cross-over experiment (Fig 3A) – red bars, pathway change in mice fed AIN76A for 3 months, then switched to NWD1 for 4 days before sacrifice; blue bars, the mice then switched back to AIN76A for 4 more days; **F**, scATACseq data for *Cyp4a10* in enterocytes; **G,** scRNAseq data for *Trpv6* Bmi1+ derived cells from AIN and NWD1 fed mice 3 days or 66-70 days after cells were marked (Fig 4A); **H**, scATACseq data for *Trpv6* in cells in enterocytes for mice fed either AIN76A or NWD1 for 4 months from weaning .

The pathway most enriched in Bmi1+ marked EC7 cells accumulating longer term was antigen processing and presentation (**Fig 6B**; NES 1.75, *P=0.009*). CD74, a key gene in this pathway, is expressed in epithelial cells (54–63), mediating interaction of Lgr5^hi^ ISCs and their immune environment (63). The fraction of cells expressing CD74 was highly enriched across all cell clusters in longer term marked Bmi1+ cells that accumulated with time (**Fig 6C**), paralleled by elevated expression per cell, with much higher expression in the EC7 mature enterocytes compared to all other clusters (**Fig 6D**, *P=10^-25^*, by negative binomial regression analysis). CD74 interacts with proteins of the major histo-compatability II (MHCII) complex to process and present cell surface antigens (58), and MHCII complex genes were also upregulated most significantly in EC7 cells (**Fig S7A**). The greatest expression of the CD74 and MHCII complex genes in EC7 cells is consistent with expression of these genes predominantly in cells in the upper third of villi (64), suggesting a unique role in these differentiated cells.

Cell surface expression of CD74 characterizes dysplastic cells of the intestinal epithelium, and cells of human Crohn’s, Ulcerative, and Amebic colitis (60,65,66), chronic inflammation in each elevating CRC risk. NWD1 is also pro-inflammatory, elevating serum levels of inflammatory cytokines (26), and there was a 2-3 fold increase in cells expressing CD3, a pan T cell marker, in 3 month NWD1 compared to AIN76A mice, significant when quantified as CD3 cells/villus (*P=0.008*), or percent area/villus epithelial column expressing CD3 (*P=0.015*) (**Fig S7B,C,D)**.

The second most highly enriched pathway induced by NWD1 in EC7 cells is “Intestinal Immune Network/IgA production” (**Fig 6B**; NES=1.50, *P=0.027*). IgA is abundant in the intestinal and colonic mucosa, traversing the lamina propria to form an SigA complex interacting with the myeloid cell FcαRI receptor, which is pro-inflammatory. NWD1 also elevated F/480+ myeloid cells (*P=0.01*; **Fig. S6 E,F**), also characterizing pro-tumorigenic IBD (67).

In EC7 cells, we also identified alteration of multiple metabolic pathways, including *Ppar* signaling, cholesterol metabolism, and fat digestion/absorption (**Fig. 6B**), adaptive responses to NWD1 increased fat (25%). In the independent scRNAseq data of the rapid dietary cross-over experiment (**Fig 3A**), fatty acid metabolism was elevated in all cell clusters by the 4-day switch to NWD1 from AIN76A, and except for Goblet2 and EE2, reverted for the clusters within 4 days of switch back to AIN76A (**Fig 6E**), consistent with rapid shift of metabolic pathways (**Fig 3B**).

Another adaptive response to NWD1 was increased number of enterocytes expressing *Trpv6* in Bmi1+ marked cells, especially longer term (**Fig 6G**). *Trpv6* encodes the enterocyte brush border calcium channel mediating calcium uptake under low dietary calcium conditions (68). Therefore, elevated *Trpv6* expression explains how NWD1 mice maintain serum calcium levels (26), despite lower calcium and vitamin D_3_ in NWD1. Single cell ATACseq data were generated from the epithelial cells of replicate mice fed AIN76A or NWD1 for 4 months. These data revealed no areas of open chromatin structure for either *Cp4a10* in enterocytes, a gene common to multiple pathways of fat metabolism, nor for the *Trpv6* gene, but major opening of chromatin structure at and upstream of the promoter in replicate libraries for NWD1 fed mice (**Fig 6F, H**), demonstrating classic epigenetic transcriptional activation of these genes generate the adaptive dietary response.

*Ppargc1a* was down regulated specifically in stem cells at the crypt base, altering mitochondrial structure that suppressed canonical Lg5^hi^ ISCs, recruiting alternates cells to maintain the mucosa. The scRNAseq had established Stem 1 harbored the Lgr5^hi^ ISCs at the crypt base, consistent with subsequent trajectory analysis identifying Stem1 at initiation of maturation of all cell types/lineages (**Fig 4B**). Of the 13 clusters identified in the scATACseq data, the cluster aligning with Stem 1 was identified using the Signace *“Reference Mapping”* function (NY Genome Center) (**Fig 7A**). The scATACseq data showed the number of these cells in this stem cell cluster decreased 50% in NWD1 mice compared to AIN76A (**Fig 7B**, *P=0.009*), independently confirming NWD1 suppression of Lgr5^hi^ cells. In contrast to the data for *Cpt4a10* and *Trpv6* (**Fig. 6F,H**), the position and magnitude of open chromatin regions at the promotor of the *Ppargc1a* in the Stem 1 cluster (designated P) is similar for all mice (**Fig 7C**). However, in the region of *Ppargc1a* delineated by the boxed region, expanded in **Fig. 7D**, “a” and “b” are areas of open chromatin in AIN76A mice absent or substantially reduced in NWD1 mice. This was quantified, showing significant reduction in signal in these regions when normalized by the strong, relatively consistent signal for all samples, at the promotor (**Fig 7E**). These regions of reduced accessibility in NWD1 mice correspond to the position of a strong enhancer in the gene with a significant role in regulating gene expression in intestinal epithelial cells (**Fig 7C,D**; *Enhanceratlas 2.0*). Thus, the scATACseq data confirm NWD1 down-regulated expression of *Ppargc1a* expression, and establish this is epigenetic, but likely by a more complex mechanism than altering accessibility of sites at or upstream of the promoter.

**Figure 7:**
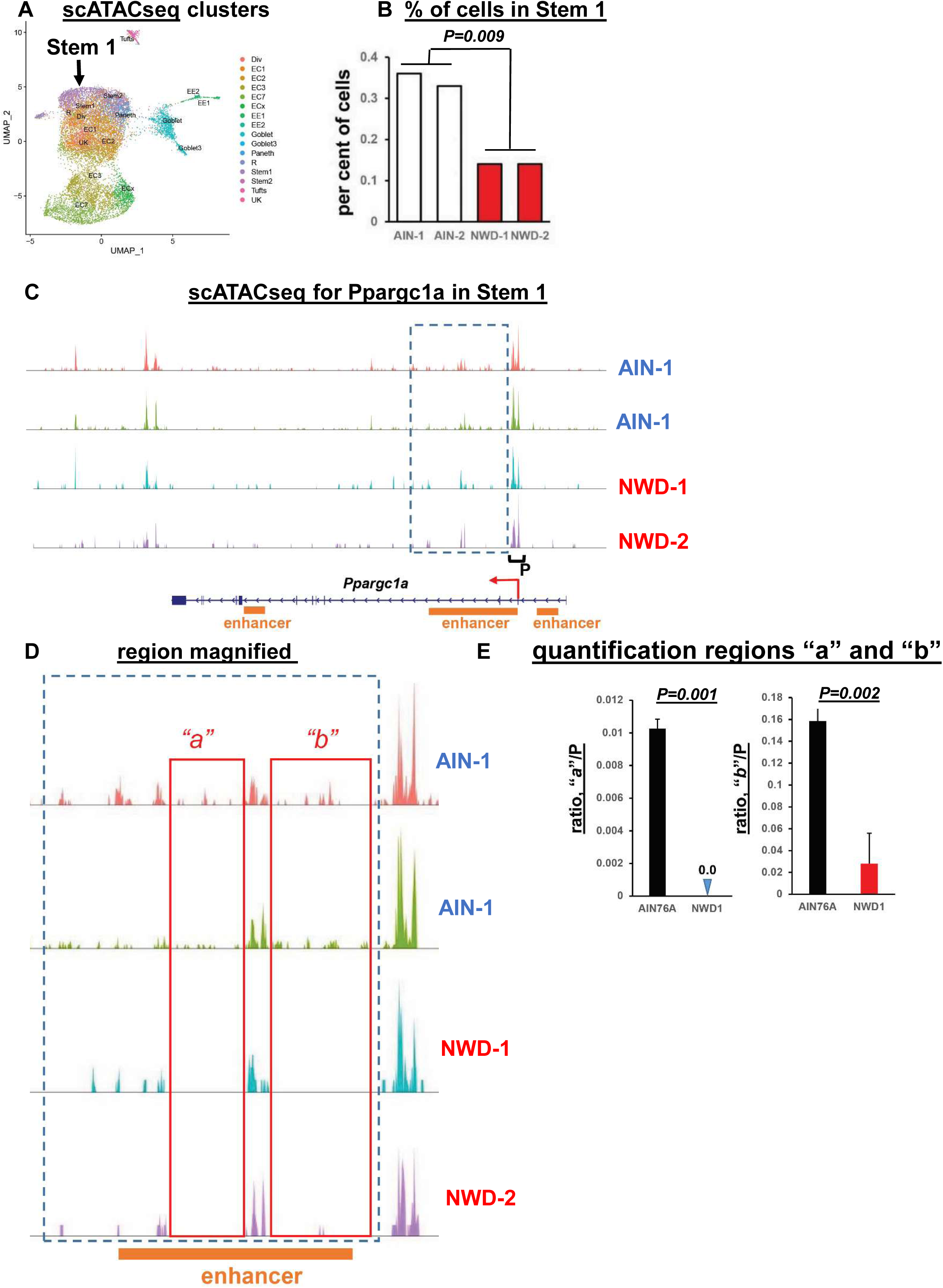
single cell ATAC seq data, stem cells: **A,** clusters from the scATACseq data; **B**, per cent Stem 1 cells in mice fed AIN76A or NWD1 based on the scATACseq data; **C,** scATACseq data for the *Ppargc1a* gene in the Stem 1 cluster. Box delineates region of diminished peaks in NWD1 compared to AIN76A fed mice, and P denotes the promoter region; **D,** boxed region in C expanded, with “a” and “b” denoting areas within this region where peaks in AIN fed mice are substantially diminished in NWD1 fed mice; **E**, quantification of reads in regions a and b relative to reads at the promotor for mice fed the different diets; statistical analysis by a poisson regression on aggregated read counts over all cells per mouse, normalized by coverage per sample in the promoter region of Ppargc1a.

## Discussion

Adjusting multiple nutrient exposures in the mouse to mimic levels elevating risk for human sporadic colon tumors causes rapid, reversible and hence dynamic adaptation of stem cells and lineages through epigenetic mechanisms that also alter pathways establishing a pro-tumorigenic environment. The data provide seven fundamental insights into dietary mechanisms of tumor risk, many paralleling pathogenic mechanisms of human inflammatory bowel disease: 1) low-level chronic inflammation that is pro-tumorigenic; 2) NWD1 reduced *Ppargc1a* expression in Lgr5^hi^ cells through epigenetic alteration of the gene, lower expression also mediating repression of Lgr5^hi^ cell stem cell functions in IBD (69), and down regulation of the encoded protein, Pgc1a, that also exacerbates human IBD (38), in both cases altering mitochondrial structure, metabolically reprogramming the cells; 3) feeding NWD1 or direct genetic inactivation of the *Ppargc1a* gene in Lgr5^hi^ cells retarded developmental maturation of cells as they migrated along the crypt-lumen axis, similar to retarded maturation of cell differentiation in human IBD along this axis also recently determined from trajectory analysis of single cell data (39); 4) NWD1 induced greatest perturbation of expression profile in mature enterocytes at the villus tip, where the most prominent change is elevation in the interacting CD74-MHCII pathways of antigen presentation and processing. This parallels highest expression of these pathways in upper villi of the human small intestine (64) and that in human IBD, upper crypt colonocytes exhibit greatest increase in this pathway (37). Further, NWD1 increased pro-inflammatory cytokines (26), and CD3 and myeloid cell mucosal infiltration, paralleling elevation of the CD74/MHCII pathways in human Crohn’s, Ulcerative, and Amebic colitis (60,65,66), confirmed by single cell analysis of human IBD tissue (37), and also in H.pylori infected gastric tissue (57, 70), all chronic inflammatory states increasing risk for human tumor development; 5) NWD1 induces high ectopic expression of Lyz and other Paneth cell markers in mouse colon (29), similar to Lyz elevation in human IBD tissue (37) and single cell RNAseq human data document expression of these markers in cells of the human mucosa that are not Paneth cells (71). Rapid alteration of Paneth cell markers in response to a 60% fat diet has also recently been reported (7). 6) In NWD1 fed mice, extensive remodeling of stem cells and lineages reflects alterations in cell reprogramming in both the involved and uninvolved mucosa in human IBD (37, 39), and in the earliest benign human tumors (72), indicating remodeling is initiated early in the mucosa at dietary risk; 7) Establishment that the nutritional environment can shift which stem cells function in mucosal maintenance parallels a recent report that human colon tumors consist of both Lgr5 positive and negative stem cells with the balance determined by the mutational signature and environmental signals from other cell types or drugs (36). The multiple parallels between NWD1 induced alterations and those in human IBD suggest that the dietary and IBD remodeling, and increased tumor risk, may share common mechanisms of pathogenesis.

The substantially increased CD74/MHCII pathways in NWD1 fed mice, similar to that in involved and uninvolved mucosa in human IBD, contrasts with an important report that a 60% fat diet repressed these pathways in mouse Lgr5^hi^ ISCs, suggested to be pro-tumorigenic by dampening immune surveillance (73). Several factors may contribute to this difference, including major differences in dietary formulation; 60% dietary fat to induce mouse obesity exceeds fat intake in even obese humans, and induces metabolic alterations differing from those induced by dietary fat levels more common in humans (15, 16). The 60% fat diet, as do all commonly used rodent diets, also includes very high levels of vitamin D exceeding the mean level in humans by 200-300%, and well above even the 1% of the population at very highest levels. This is fundamental since robust expression of the vitamin D receptor is a core component of the Lgr5^hi^ stem cell signature, and we showed and extensively discussed the essential role of vitamin D in Lgr5^hi^ stem cell function, as it is in hair follicle and other stem cells (8,9,74). Further, as discussed in detail (4,6,8,9), the much lower vitamin D levels in humans likely contributes to the fact that it takes 50 fold longer for CBC stem cells in humans to reach clonality than in the mouse (1,75,76). Thus, the 60% fat diet, and other commonly used mouse diets, establish a nutritional environment strongly supporting Lgr5^hi^ ISC functions, but this environment necessary for robust function of Lgr5^hi^ stem cells is simply not present in nearly all humans.

NWD1 remodeling of stem cells and lineages includes epigenetic regulation of pathways mediating the rapid and reversible adaptation of the mucosa to its nutritional environment. This included upregulated *Trpv6* expression in enterocytes, the gut calcium transporter maintaining calcium homeostasis under low calcium conditions, explaining how, despite lower calcium in NWD1, serum calcium level is maintained with no loss of bone mineral density (26). A second example is rapid and reversible alteration of fat metabolism pathways in response to the 25% fat in NWD1, as recently reported for a 60% fat diet (7). For both NWD1 examples, elevated expression paralleled opening of chromatin configuration at the gene promoter, independently confirming response to NWD1 and demonstrating epigenetic regulation.

The significance of the metabolic reprogramming of Lgr5^hi^ stem cells by NWD1 is emphasized by necessity of Oxphos for mouse Lgr5^hi^ and Drosophila intestinal stem cells to function efficiently as stem cells (40–42), for hematopoietic stem cell functions (43), for embryonic stem cell pluripotency (44), and a major role of mitochondrial function in determining whether stem cells self-renew or differentiate (51). The repression of Lgr5^hi^ cells as stem cells recruits alternate *Bmi1+, Ascl2^hi^* cells above the crypt base for mucosal maintenance, as in response to acute damage. *Ascl2* encodes a stem cell transcription factor essential for Bmi1+ cell de-differentiation in response to Lgr5^hi^ cell damage (1, 2). *Ascl2* is regulated by Wnt signaling (77), elevated by NWD1 throughout mouse intestinal villi and colonic cells (27–29). However, there are likely different roles of *Bmi1+, Ascl2^hi^* cells in NWD1 versus damage induced plasticity. Acute damage purges Lgr5^hi^ ISCs mobilizing *Ascl2^hi^* cells to migrate into the crypt to restore crypt organization. However, limited crypt space establishes competition among crypt Lgr5^hi^ cells (1), and single cell laser ablation of crypt base cells triggers division and reorganization of remaining stem cells, confirming importance of physical space in regulating stem cell dynamics (78). In contrast, crypt space may be unavailable with chronic NWD1 feeding. Further, cells migrating into the niche would still be repressed by NWD1, and reduced Lgr5^hi^ cell lineage tracing from the crypt base, and Oxphos and TCA pathway repression, persist to at least 1 year of feeding NWD1 (8) (**Fig S1, 2**), with Bmi1+ cells above the crypt base lineage tracing for months in mice continuously fed NWD1 (6).

Tumors reflect growth advantage of transformed cells (1), but this is very rare: lower-risk individuals develop no sporadic tumors, and those at higher risk only 1-2 tumors despite billions of mucosal cell divisions over decades, suggesting mechanisms establishing risk are subtle and complex. Therefore, it is fundamental that the nutritional environment, through metabolic reprogramming of stem cells, dynamically and continually sculpts the playing field on which intestinal stem cell competition takes place, with significant influence in determining outcome.

## Experimental Procedures

### Mice

Mice, on a C57Bl/6J background, were provided food and water *ad libitum* in a barrier facility at the Albert Einstein College of Medicine. For breeding, strains were fed a chow diet (Picolab 5058, Fisher Feed, Branchburg, NJ). Appropriate genotypes were randomized to purified diets (AIN76A, D10011; NWD1, D16378C; Research Diets Inc, New Brunswick, NJ). Mice of both genders were used (a total of 25 male and 20 female mice). Tamoxifen in corn oil was a single injection (Sigma, T5648, 100µl, 1mg/µl). Experiments were approved by the Albert Einstein Institutional Animal Care and Use Committee. All authors had access to the study data.

### Isolation of intestinal epithelial cells

On sacrifice, excised intestines were opened longitudinally, rinsed with cold saline, crypts isolated, single cell suspensions prepared, and depending on the experiment stained with EpCAM-FITC+ (Miltenyi Biotec, Clone caa7-9G8, Cat No. 130-123-674), CD45-PerCP negative (Miltenyi Biotec, Clone 30F11, Cat No. 130-123-879) sorted by FACs and collected on a MoFlo instrument as described (6)

### Bulk RNAseq analysis

construction of bulk RNAseq libraries, sequencing and data analysis were as described (6).

### Single cell RNAseq

Single cell RNAseq (scRNAseq) libraries were constructed by the Albert Einstein Genomics Core using the 3’ kit protocols of 10X Genomics (San Francisco, CA) with approximately 10,000 single FACS collected cells from each mouse loaded onto a Chromium Chip B microfluidic apparatus. Library quality was verified by size analysis (BioAnalyzer; Agilent, Santa Clara, CA) and after passing quality control assays, multiple libraries mixed and sequenced by HiSeq PE150 or NovaSeq PE150 using pair-end sequencing with a readout length of 150 bases (Novogene; Sacramento, California), and the data assigned to individual cells using the sample indexes.

For sequence alignment and subsequent analysis, output Illumina base call files were converted to FASTQ files (bcl2fastq), aligned to the mouse mm10 genome v1.2.0 and converted to count matrices (Cell Ranger software v3.0.2). Force-cell parameter was used, with 5000-8000 individual cells identified for each sample, and unique molecular identifiers (UMI) identified to remove PCR duplicates. Quality control and downstream analyses were done in R v4.0.2, using Seurat package v3.2.2 (79). To discard doublets or dead cells, cells with gene counts <200 or >5000, or a mitochondrial gene ratio >20%, were filtered out. Cells from different samples were integrated using Seurat FindIntegrationAnchors and IntegrateData functions and clusters identified from each integrated data set using Seurat FindClusters. This calculates k-nearest neighbors by Principle Component Analysis, constructs a shared nearest neighbor graph, and then optimizes a modularity function (Louvain algorithm) to determine cell clusters. Cell clusters were identified as cell intestinal cell types using established cell-type markers and cluster identification using Seurat FindMarkers function. Differential gene expression compared samples among experimental groups: initial criteria were an expression change of ≥1.5 fold with associated adjusted P-value of <0.01 in group comparison (Seurat FindMarkers function). Pathway analysis was performed on differentially expressed genes using clusterProfiler R package v3.16.1 and the KEGG pathway database (KEGG.db R package v3.2.4); gene set enrichment analysis (GSEA) used the fgsea R package v1.14.0, the MSigDB (v5.2) hallmark gene set and the KEGG pathway database. Pathways at *P <0.05* were considered statistically significant. Trajectory analysis used Monocle3 R package v0.2.3.3 (80) with cluster information and cell-type identifications migrated from Seurat, the integrated dataset divided among the experimental conditions, and trajectories generated for experimental group.

RNAseq data have been deposited in the NCBI Gene Expression Omnibus (GEO) database: bulk RNAseq (Fig 1) - GSE186811; scRNAseq (Fig 2, *Ppargc1a* inactivation) - GSE188339; scRNAseq (Fig 3, rapid dietary crossover) - GSE188577; scRNAseq for Bmi1+ cells (Fig 4.5) - GSE188338.

### *In situ* analyses

Sections of mouse intestinal swiss rolls fixed in 10% formalin were de-waxed and rehydrated. Reagents were from Advanced Cell Diagnostics (Newark, CA). Endogenous peroxidases were quenched and heat induced epitope retrieval done utilizing the RNAscope Multiplex Fluorescent Reagent Kit v2 (Cat. No. 323100) and the RNAscope Multiplex Fluorescent Assay v2 protocol followed. RNAscope target probes for *Bmi1* and *Ascl2* Cat No. 466021, 412211, respectively) were hybridized to sections and amplified using the HybEZ Hybridization System (Cat No. 310010). Secondary TSA fluorophores from Akoya Biosciences were Opal 620 (channel 1, Cat No. FP1495001KT) and Opal 570 (channel 2, FP1488001KT), sections counterstained with Dapi and visualized using a Leica SP8 confocal microscope (20X magnification).

### Single cell ATACseq

Total Epcam+, CD45 negative epithelial cells from the small intestine of mice were FACs isolated as for scRNAseq. Nuclei were isolated following the 10X Genomics protocol, single cell ATACseq libraries constructed using the 10X Geomics single cell ATACseq Chromium reagents and protocol, and sequenced as for scRNAseq.

### Immunohistochemistry

Swiss roll sections were de-waxed, rehydrated, endogenous peroxidases blocked (3% hydrogen peroxide in methanol) and heat induced epitope retrieval done with 10 mM Sodium citrate buffer (pH 6.0) for all antibodies. Tissues were incubated overnight at 4^0^C with primary CD3 antibody (Novusbio, Littleton, CO, 1:10 dilution of NB600-1441). Protein block and secondary antibody for CD3 employed ImmPRESS-Alkaline Phosphatase Horse Anti-Rabbit IgG Polymer Kit-Alkaline Phosphatase (Vector Laboratories, Burlingame, CA, Cat No. MP-5401). F4/80 antibody (1:200, Cell Signaling, Denvers, MA, Cat No. 70076, Cell Signaling, Denvers, MA) was incubated on tissue overnight at 4^0^C. Secondary antibody was SignalStain® Boost IHC Detection Reagent (HRP, Rabbit, Cell Signaling, Cat No. 8114). For Pgc1a staining, tissues were incubated overnight at 4 °C overnight with rabbit anti-Pgc1a Ab (diluted 1:500, NBP1-04676, Novusbio) followed by biotinylated goat anti-rabbit IgG (1:100, BA-1000; Vector Lab,) for 30 min. Visualization used the Vectastain Elite ABC Kit (Vector Lab) and DAB Quanto kit (Thermo Fisher Scientific). Counterstaining was with hematoxylin and visualization by bright field microscopy, or a CY5 filter for CD3.

### Electron Microscopy

Samples were fixed with 2.5% glutaraldehyde, 2% paraformaldehyde in 0.1 M sodium cacodylate buffer, post-fixed with 1% osmium tetroxide followed by 2% uranyl acetate, dehydrated through a graded ethanol series and embedded in LX112 resin (LADD Research Industries, Burlington VT). Ultrathin sections were cut on a Leica Ultracut UC7, stained with uranyl acetate followed by lead citrate and viewed on a JEOL 1400 Plus transmission electron microscope at 120kv. Images were captured at a magnification of 800X, and at 5000X for quantitative measurements of mitochondrial size and cristae density.

### Mitochondrial membrane potential

Crypts were dispersed into single-cell suspensions, and cells incubated with MitoTracker^TM^ Red FM (Invitrogen, Cat # M22425, 25nM) for 20 min at 37 °C. Zombie NIR (Invitrogen, Cat # L10119) was then added to mark dead cells. Flow cytometry used a Cytek Aurora Instrument (Cytek Biosciences) with analysis using Flowjo 10 software (Treestar Inc.).

### Lineage tracing of *Ppargc1a* knock-out mice

Induction of *cre^ERT2^*activity was by a 100 μl intraperitoneal injection of 1 mg freshly prepared Tamoxifen (TAM) in sterile corn oil. Mice were sacrificed 3 days after injection, intestinal tissues fixed with 4% paraformaldehyde for 2 hours, hydrated with 15% sucrose for 3 hours, and 30% sucrose overnight. Fixed tissues were embedded in optimal cutting temperature (OCT) compound (Sakura), frozen on dry ice, cryostat sectioned, and stained with DAPI for identification of nuclei before imaging.

### Image analysis

To quantify anti-Pgc1a Ab staining, images were at 60x magnification. A threshold was applied for all slides, and the area above threshold calculated and divided by total area. The crypt-villus axis was divided by 20 μm increments from the base of the crypt and measured up to 80 μm. For analysis of mitochondrial morphometry, crypt base columnar cells, Paneth cells, and villus cells imaged at 5000x magnification were selected. Cristae density was calculated by the sum of individual cristae divided by mitochondrial size. Mitochondrial size (μm^2^) and cristae area (μm^2^) were manually traced using Fiji. For lineage tracing quantification, the crypt-villus axis was manually traced and the threshold automatically adjusted for each crypt. Red fluorescent positive area (μm^2^) was measured and divided by total crypt-villus area (μm^2^).

## Acknowledgements

Supported in part by R01CA229216, R01CA214625, R01CA222358 and P30CA013330 from the NCI, NIH, USPHS.

---

**Figure S1:**
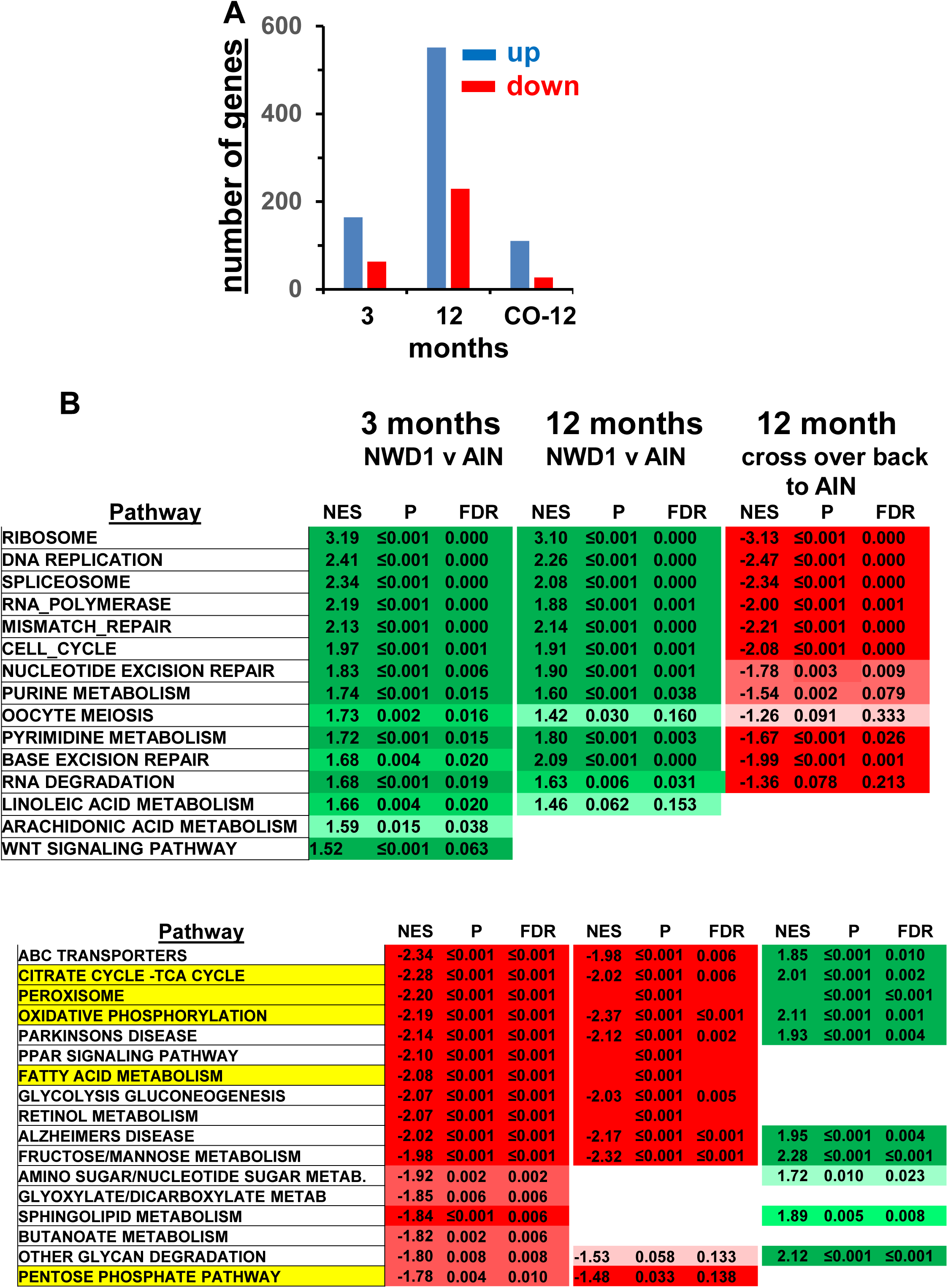
**A,** number of the ∼8000 expressed sequences in Lgr5^hi^ cells up or down-regulated (≥50%, *P≤0.01)* at 3 or 12 months feeding NWD1 compared to AIN76A, or 3 months NWD1 then switched to AIN76A for 9 months (Arm 3, Fig 1); **B,** Pathways significantly altered by Gene Set Enrichment Analysis of bulk RNAseq data of Lgr5^hi^ cells isolated by FACS from mice fed NWD1 compared to AIN76A for either 3 or 12 months, or for mice fed NWD1 for 3 months than switched back to AIN76A for 9 months, described in Fig 1A.

**Figure S2:**
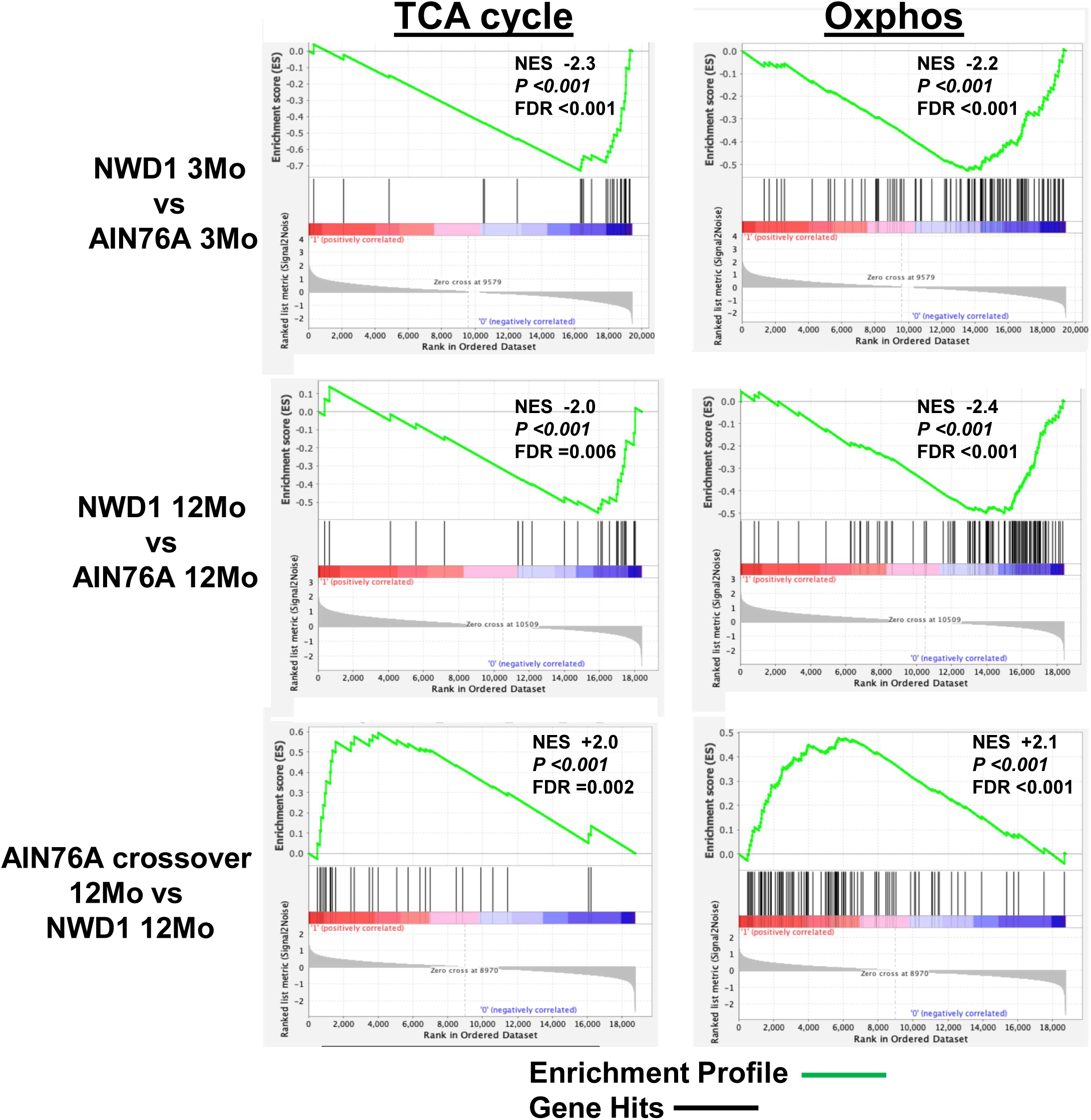
GSEA of the TCA cycle and Oxphos pathways in mice of the 3 Arms of Fig. 1A. Inserts in each panel show the NES, *P-value* and false discovery rate (FDR).

**Figure S3:**
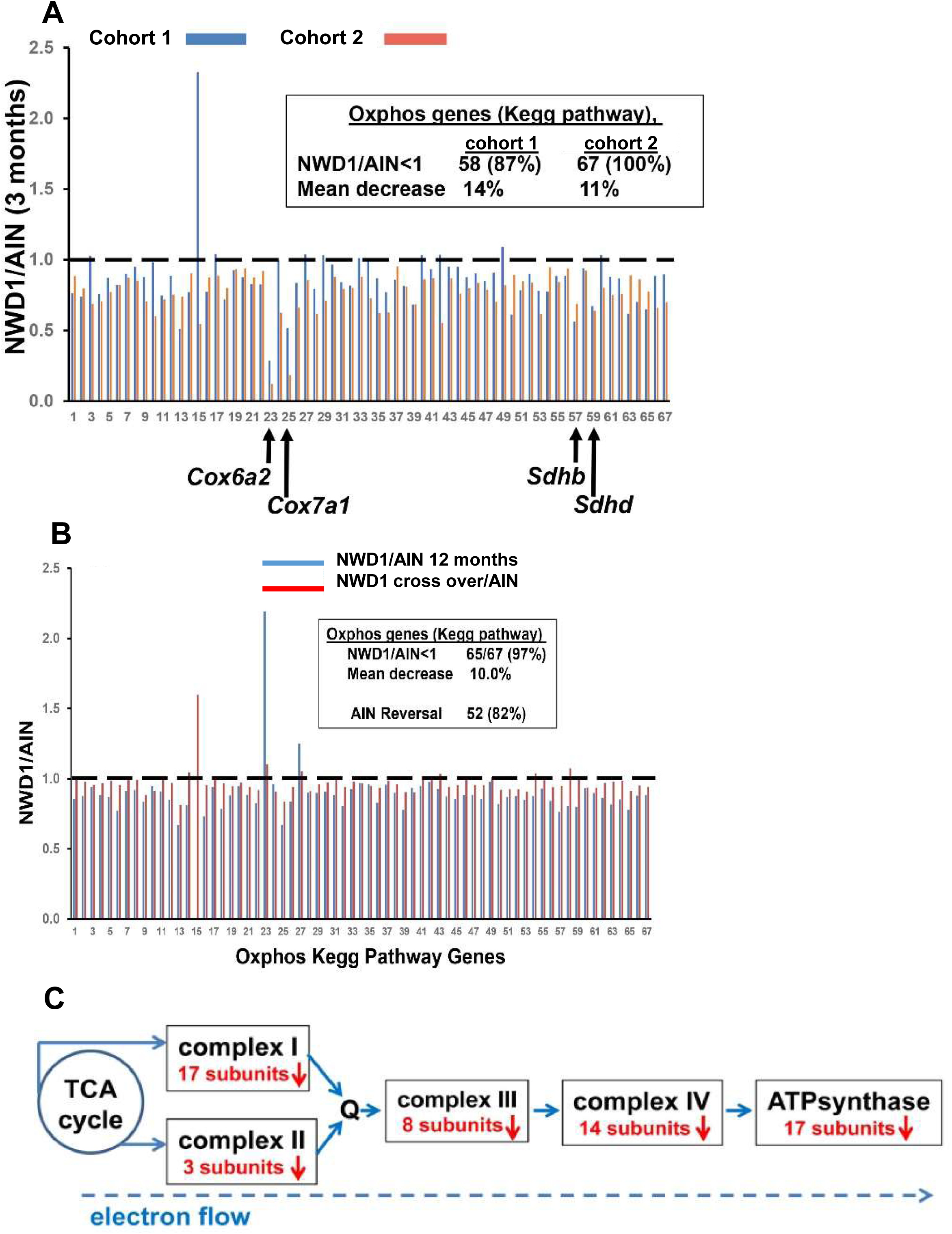
**A,** Expression ratio (bulk RNAseq) of Oxphos pathway genes in Lgr5^hi^ cells of mice fed NWD1 vs AIN76A for 3 months (Fig 1A) or of an independent bulk RNAseq data set from different mouse cohorts (6) and **B**, Bulk RNAseq expression ratio of Oxphos pathways genes in Lgr5^hi^ cells fed NWD1 compared to AIN76A for 12 months or in the cross over arm of Fig 1A, NWD1 for 3 months switched to AIN76A for an additional 9 months (12 months total) compared to mice fed NWD1 continuously for 12 months. Statistical analysis detailed in text; **C**, summary of the down-regulated genes encoding subunits of each of the 5 mitochondrial electron transport chain complexes.

**Figure S4:**
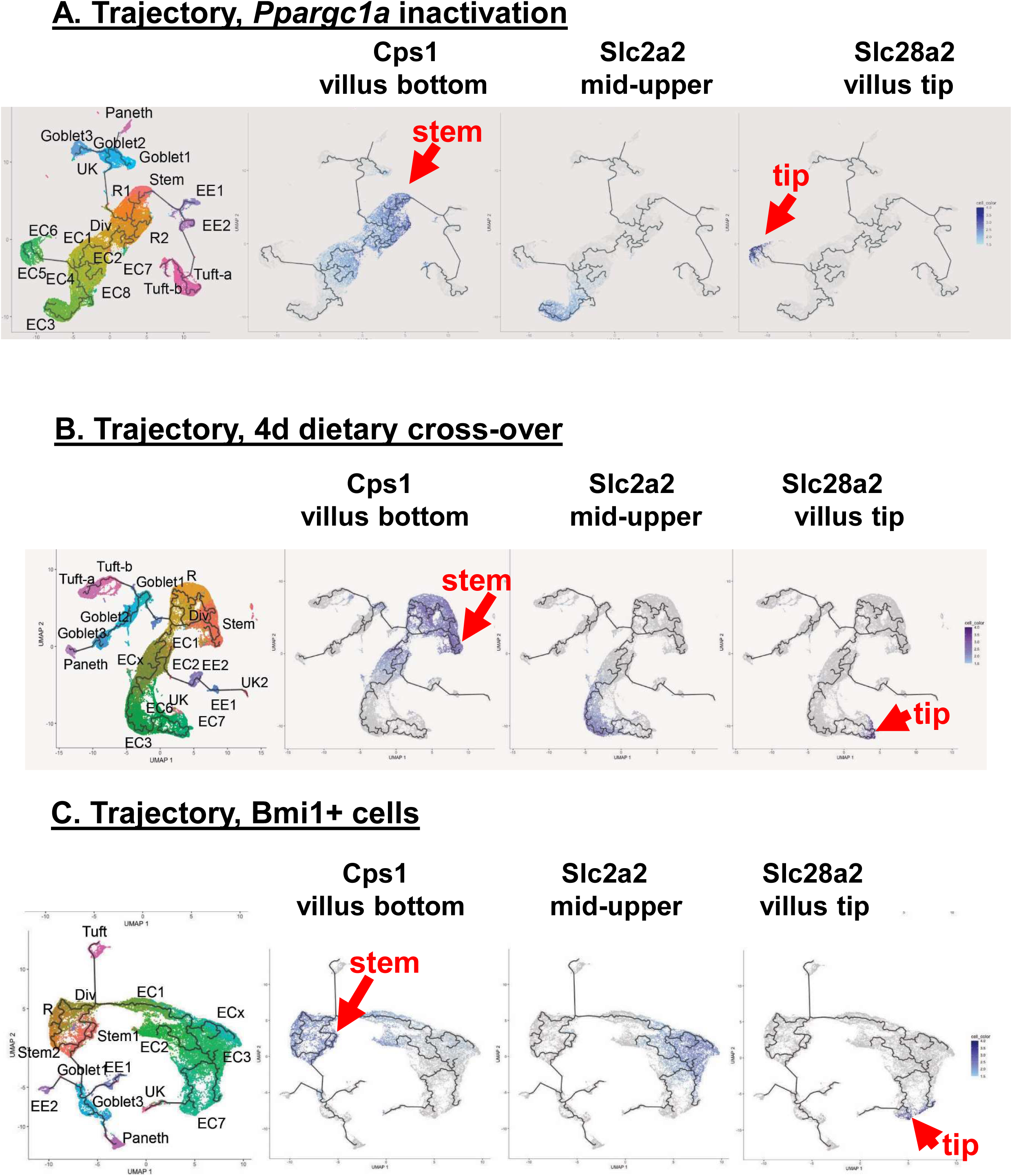
Validation of predicted trajectory from scRNAseq data by expression of markers shown to be physically localized along the crypt-villus axis (villus bottom, mid-upper villus and villus tip) (50). **A**, trajectories of Fig 2, *Ppargc1a* genetic inactivation in Lgr5^hi^ cells; **B**, trajectories of Fig 3, rapid dietary cross-over; **C**, trajectories of Fig 4A and 5A, Bmi1+ marked cells.

**Fig S5:**
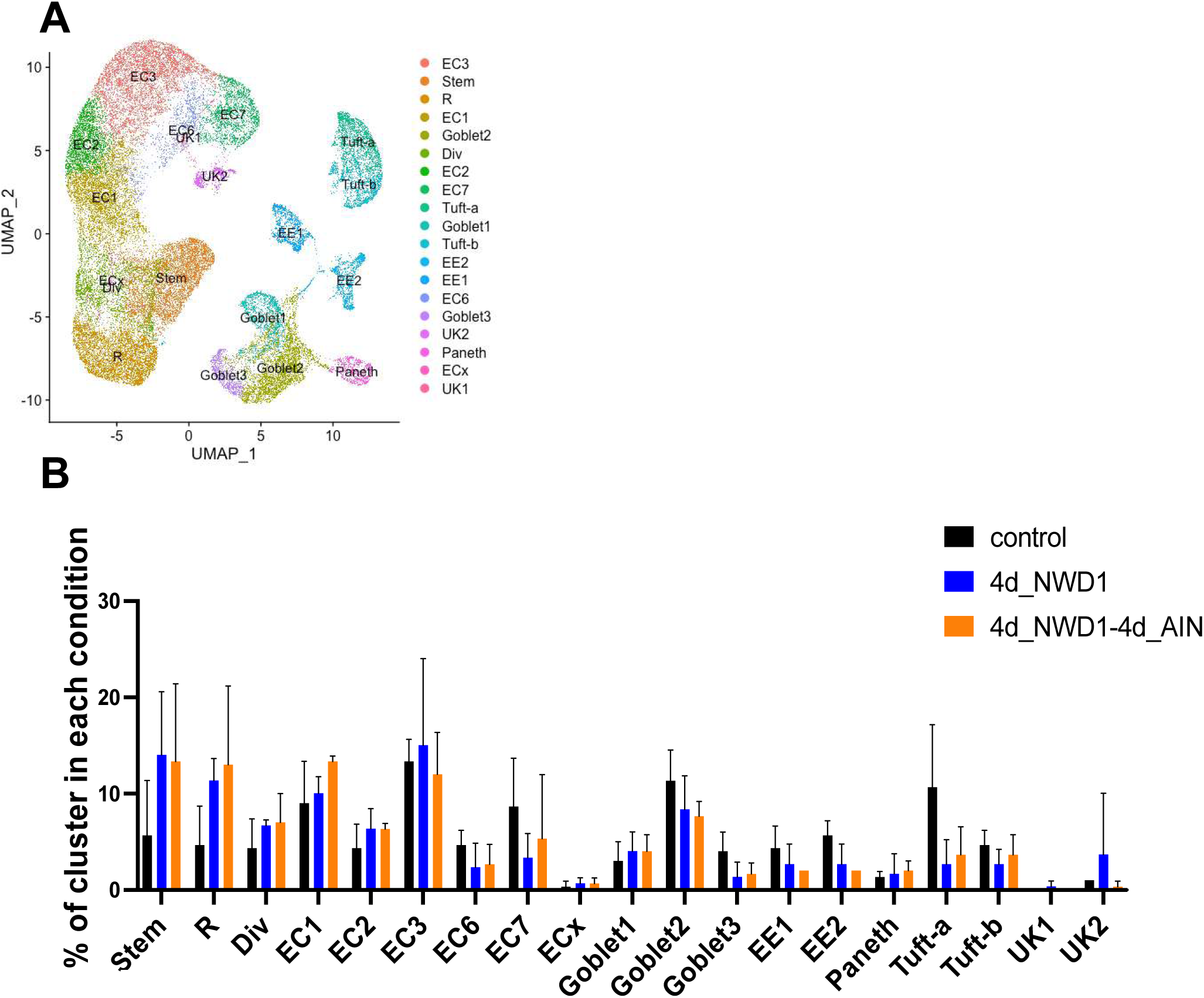
Rapid dietary cross-over experiment of Fig 3. **A,** cluster map and cell types, N=3 independent mice for each dietary Arm (9 scRNAseq libraries – Fig 3A); **B**, epithelial cell type distribution in clusters/lineages.

**Fig S6:**
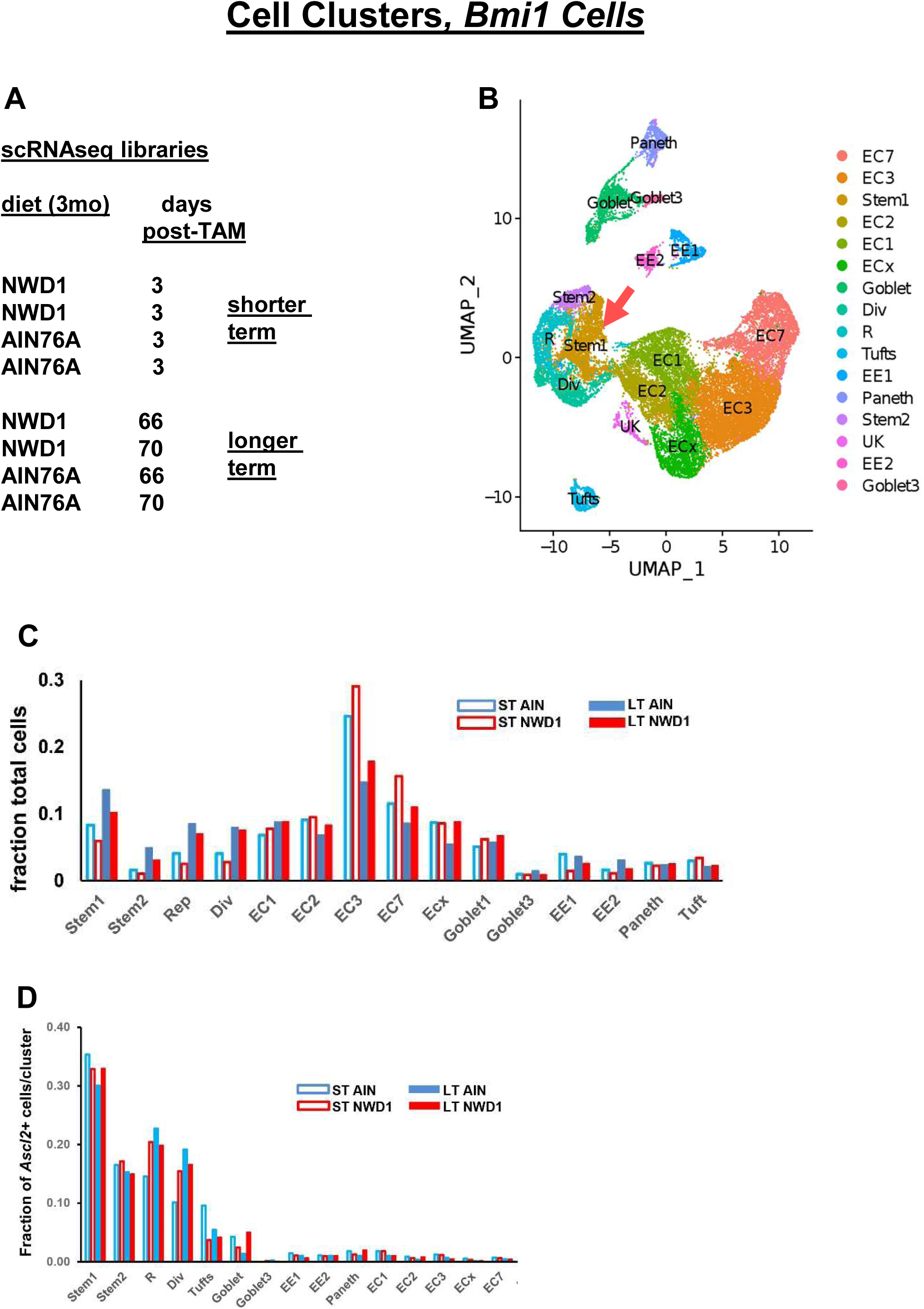
scRNAseq analysis of Bmi1+ derived cells. **A**, Bmi1+ derived epithelial cells from mice fed AIN76A or NWD1 FACs isolated 3 or 66-70 days post marking (Fig 5); **B,** cluster map of intestinal epithelial cell types/lineages – red arrow, stem cell cluster**; C,** intestinal epithelial cell type/lineage distribution by diet and time post Tam marking of cells; **D,** fraction of cells Ascl2 positive in each cluster by diet and time post marking.

**Fig S7:**
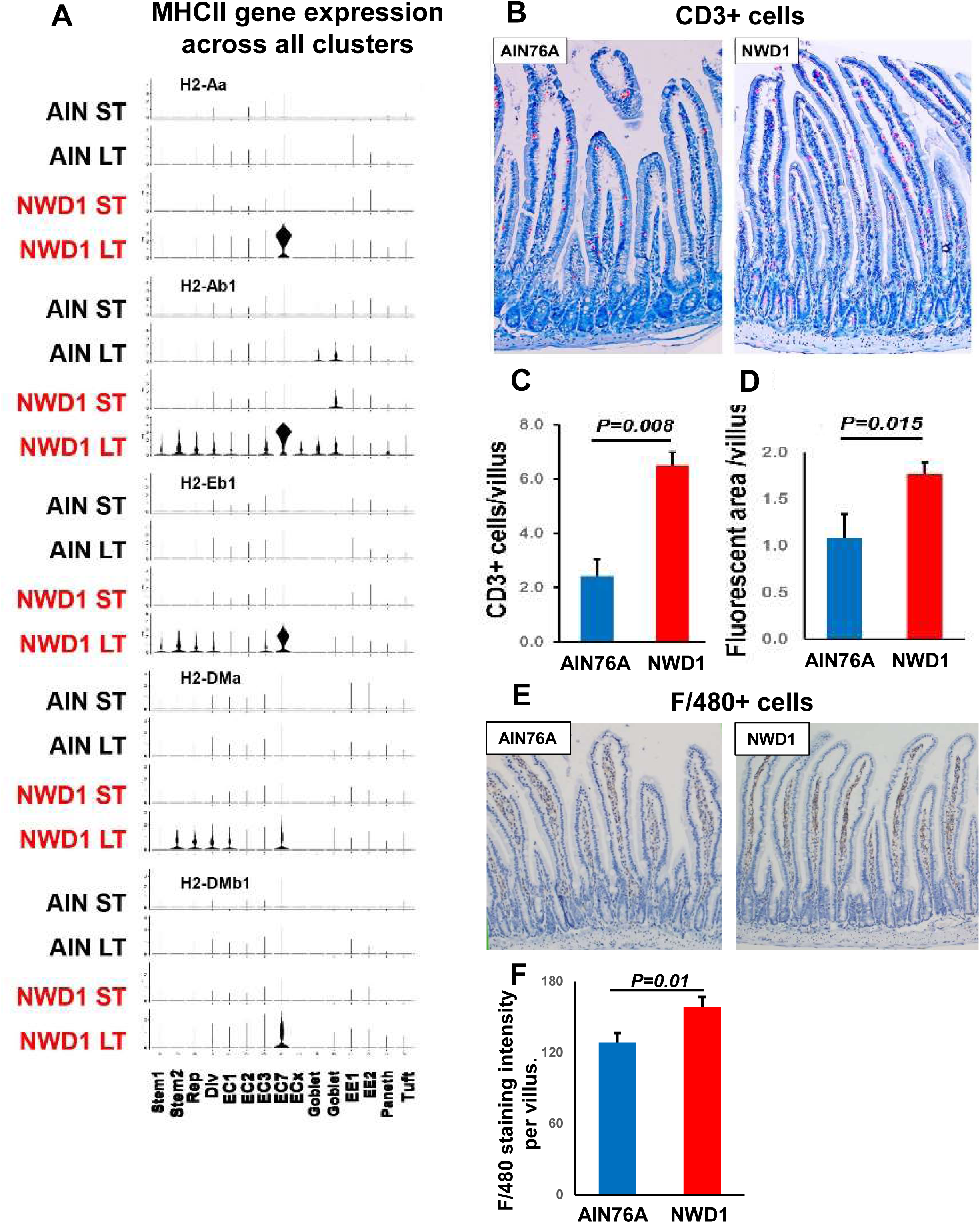
Inflammatory response to NWD1: **A,** expression of 5 genes encoding proteins of the MHCII pathway across cell type/lineages in relation to diet and shorter term (ST= 3d) or longer term (LT=66-70d) post Tam marking of Bmi1+ cells; **B**, immunohistochemical detection of CD3+ cells in AIN76A and NWD1 fed mice; **C, D,** quantitation of CD3+ cells in relation to diet by number of CD3+/villus (**C**) or area of red fluorescent staining in the mucosa (**D**), N=3 mice per dietary group; **E**, F480+ cells in AIN76A and NWD1 fed mice; **F**, staining intensity per villus, N=3 mice per dietary group.

