## Supplementary material for "Dynamic intestinal stem cell plasticity and lineage remodeling by a nutritional environment relevant to human risk for tumorigenesis": Table S1 (from another version of this preprint: version 2, 2022-08-15)

### Table S1: All pathways significantly altered in $Lgr5^{hi}$ cells of NWD1 fed mice

| Pathway | 3 months | | | 12 months | | | NWD1/AIN cross-over | | |
|---|---|---|---|---|---|---|---|---|---|
| | NES | P value | FDR | NES | P value | FDR | NES | P value | FDR |
| _RIBOSOME | 3.190 | ≤0.001 | 0.000 | 3.100 | ≤0.001 | 0.000 | -3.130 | ≤0.001 | 0.000 |
| _DNA_REPLICATION | 2.410 | ≤0.001 | 0.000 | 2.260 | ≤0.001 | 0.000 | -2.470 | ≤0.001 | 0.000 |
| _SPLICEOSOME | 2.340 | ≤0.001 | 0.000 | 2.080 | ≤0.001 | 0.000 | -2.340 | ≤0.001 | 0.000 |
| _RNA_POLYMERASE | 2.190 | ≤0.001 | 0.000 | 1.880 | ≤0.001 | 0.001 | -2.000 | ≤0.001 | 0.001 |
| _MISMATCH_REPAIR | 2.130 | ≤0.001 | 0.000 | 2.140 | ≤0.001 | 0.000 | -2.210 | ≤0.001 | 0.000 |
| _CELL_CYCLE | 1.970 | ≤0.001 | 0.001 | 1.910 | ≤0.001 | 0.001 | -2.080 | ≤0.001 | 0.000 |
| _NUCLEOTIDE_EXCISION_REPAIR | 1.830 | ≤0.001 | 0.006 | 1.900 | ≤0.001 | 0.001 | -1.780 | 0.003 | 0.009 |
| _TASTE_TRANSDUCTION | 1.780 | ≤0.001 | 0.011 | | | | | | |
| _PURINE_METABOLISM | 1.740 | ≤0.001 | 0.015 | 1.600 | ≤0.001 | 0.038 | -1.540 | 0.002 | 0.079 |
| _OOCYTE_MEIOSIS | 1.730 | 0.002 | 0.016 | 1.420 | 0.030 | 0.160 | -1.260 | 0.091 | 0.333 |
| _PYRIMIDINE_METABOLISM | 1.720 | ≤0.001 | 0.015 | 1.800 | ≤0.001 | 0.003 | -1.670 | ≤0.001 | 0.026 |
| _BASE_EXCISION_REPAIR | 1.680 | 0.004 | 0.020 | 2.090 | ≤0.001 | 0.000 | -1.990 | ≤0.001 | 0.001 |
| _RNA_DEGRADATION | 1.680 | ≤0.001 | 0.019 | 1.630 | 0.006 | 0.031 | -1.360 | 0.078 | 0.213 |
| _LINOLEIC_ACID_METABOLISM | 1.660 | 0.004 | 0.020 | 1.460 | 0.062 | 0.153 | | | |
| _ARACHIDONIC_ACID_METABOLISM | 1.590 | 0.015 | 0.038 | | | | | | |
| _WNT_SIGNALING_PATHWAY | 1.520 | ≤0.001 | 0.063 | | | | | | |
| _ALPHA_LINOLENIC_ACID_METABOLISM | 1.490 | 0.042 | 0.072 | 1.400 | 0.068 | 0.189 | | | |
| _DORSO_VENTRAL_AXIS_FORMATION | 1.470 | 0.047 | 0.082 | | | | | | |
| _PROGESTERONE_MEDIATED_OOCYTE_MATURATION | 1.450 | 0.021 | 0.088 | 1.440 | 0.018 | 0.157 | | | |
| _ETHER_LIPID_METABOLISM | 1.410 | 0.072 | 0.116 | | | | | | |
| _AMINOACYL_TRNA_BIOSYNTHESIS | | | | 1.440 | 0.047 | 0.149 | -1.530 | 0.031 | 0.073 |
| _HOMOLOGOUS_RECOMBINATION | | | | 1.960 | ≤0.001 | 0.001 | -2.430 | ≤0.001 | 0.000 |
| _BASAL_CELL_CARCINOMA | | | | 1.460 | 0.034 | 0.160 | -1.360 | 0.078 | 0.217 |
| _NOTCH_SIGNALING_PATHWAY | | | | 1.450 | 0.030 | 0.160 | | | |
| _BASAL_TRANSCRIPTION_FACTORS | | | | 1.390 | 0.072 | 0.187 | -1.210 | 0.209 | 0.388 |

| Pathway | NES | P value | FDR | NES | P value | FDR | NES | P value | FDR |
|---|---|---|---|---|---|---|---|---|---|
| _ABC_TRANSPORTERS | -2.340 | ≤0.001 | ≤0.001 | -1.980 | ≤0.001 | 0.006 | 1.850 | ≤0.001 | 0.010 |
| _CITRATE_CYCLE_TCA_CYCLE | -2.280 | ≤0.001 | ≤0.001 | -2.020 | ≤0.001 | 0.006 | 2.010 | ≤0.001 | 0.002 |
| _PEROXISOME | -2.200 | ≤0.001 | ≤0.001 | | ≤0.001 | | | ≤0.001 | ≤0.001 |
| _OXIDATIVE_PHOSPHORYLATION | -2.190 | ≤0.001 | ≤0.001 | -2.370 | ≤0.001 | ≤0.001 | 2.110 | ≤0.001 | 0.001 |
| _PARKINSONS_DISEASE | -2.140 | ≤0.001 | ≤0.001 | -2.120 | ≤0.001 | 0.002 | 1.930 | ≤0.001 | 0.004 |
| _PPAR_SIGNALING_PATHWAY | -2.100 | ≤0.001 | ≤0.001 | | ≤0.001 | | | | |
| _FATTY_ACID_METABOLISM | -2.080 | ≤0.001 | ≤0.001 | | ≤0.001 | | | | |
| _GLYCOLYSIS_GLUCONEOGENESIS | -2.070 | ≤0.001 | ≤0.001 | -2.030 | ≤0.001 | 0.005 | | | |
| _RETINOL_METABOLISM | -2.070 | ≤0.001 | ≤0.001 | | ≤0.001 | | | | |
| _ALZHEIMERS_DISEASE | -2.020 | ≤0.001 | ≤0.001 | -2.170 | ≤0.001 | ≤0.001 | 1.950 | ≤0.001 | 0.004 |
| _FRUCTOSE_AND_MANNOSE_METABOLISM | -1.980 | ≤0.001 | ≤0.001 | -2.320 | ≤0.001 | ≤0.001 | 2.280 | ≤0.001 | ≤0.001 |
| _ENDOCYTOSIS | -1.980 | ≤0.001 | ≤0.001 | | | | | | |
| _AMINO_SUGAR_AND_NUCLEOTIDE_SUGAR_METABOLISM | -1.920 | 0.002 | 0.002 | | | | 1.720 | 0.010 | 0.023 |
| _GLYOXYLATE_AND_DICARBOXYLATE_METABOLISM | -1.850 | 0.006 | 0.006 | | | | | | |
| _INTESTINAL_IMMUNE_NETWORK_FOR_IGA_PRODUCTION | -1.840 | 0.006 | 0.006 | -1.520 | 0.049 | 0.123 | | | |
| _SPHINGOLIPID_METABOLISM | -1.840 | ≤0.001 | 0.006 | | | | 1.890 | 0.005 | 0.008 |
| _BUTANOATE_METABOLISM | -1.820 | 0.002 | 0.006 | | | | | | |
| _EPITHELIAL_CELL_SIGNALING_IN_HELICOBACTER_PYLORI_INFECTION | -1.820 | ≤0.001 | 0.007 | -1.650 | ≤0.001 | 0.070 | | | |
| _OTHER_GLYCAN_DEGRADATION | -1.800 | 0.008 | 0.008 | -1.530 | 0.058 | 0.133 | 2.120 | ≤0.001 | ≤0.001 |
| _PENTOSE_PHOSPHATE_PATHWAY | -1.780 | 0.004 | 0.010 | -1.480 | 0.033 | 0.138 | | | |
| _CARDIAC_MUSCLE_CONTRACTION | | | | -1.800 | ≤0.001 | 0.028 | | | |
| _HUNTINGTONS_DISEASE | | | | -1.520 | ≤0.001 | 0.127 | 1.290 | 0.049 | 0.203 |
| _SYSTEMIC_LUPUS_ERYTHEMATOSUS | | | | | ≤0.001 | | | | |
| _STEROID_HORMONE_BIOSYNTHESIS | | | | -2.060 | ≤0.001 | 0.004 | | | |
| _TRYPTOPHAN_METABOLISM | | | | -2.010 | ≤0.001 | 0.005 | | | |
| _RIG_I_LIKE_RECEPTOR_SIGNALING | | | | -1.670 | 0.012 | 0.069 | 1.870 | ≤0.001 | 0.009 |
| _DRUG_METABOLISM_CYTOCHROME_P450 | | | | -1.580 | 0.011 | 0.109 | 1.330 | 0.086 | 0.179 |
| _GALACTOSE_METABOLISM | | | | -1.560 | 0.051 | 0.114 | 1.550 | 0.029 | 0.067 |
| _METABOLISM_XENOBIOTICS_BY_CYT_P450 | | | | -1.500 | 0.031 | 0.131 | | | |
| _PHOSPHATIDYLINOSITOL_SIGNALING | | | | -1.410 | 0.026 | 0.202 | 1.740 | 0.005 | 0.020 |

**Table S1: Pathways significantly altered:** GSEA of bulk RNAseq analysis of $Lgr5^{hi}$ cells isolated by FACS from mice fed NWD1 compared to AIN76A for either 3 or 12 months, or for mice fed NWD1 for 3 months than switched back to AIN76A for 9 months, described in Fig 1A.
